# Differential regulation of mRNA fate by the human Ccr4-Not complex is driven by CDS composition and mRNA localisation

**DOI:** 10.1101/2021.03.13.435226

**Authors:** Sarah L. Gillen, Chiara Giacomelli, Kelly Hodge, Sara Zanivan, Martin Bushell, Ania Wilczynska

## Abstract

**Background:** Regulation of protein output at the level of translation allows for a rapid adaptation to dynamic changes to the cell’s requirements. This precise control of gene expression is achieved by complex and interlinked biochemical processes that modulate both the protein synthesis rate and stability of each individual mRNA. A major factor coordinating this regulation is the Ccr4-Not complex. Despite playing a role in most stages of the mRNA life cycle, no attempt has been made to take a global integrated view of how the Ccr4-Not complex affects gene expression.

**Results:** This study has taken a comprehensive approach to investigate post-transcriptional regulation mediated by the Ccr4-Not complex assessing steady-state mRNA levels, ribosome position, mRNA stability and protein production transcriptome-wide. Depletion of the scaffold protein CNOT1 results in a global upregulation of mRNA stability and the preferential stabilisation of mRNAs enriched for G/C-ending codons. We also uncover that mRNAs targeted to the ER for their translation have reduced translational efficiency when CNOT1 is depleted, specifically downstream of the signal sequence cleavage site. In contrast, translationally upregulated mRNAs are normally localised in p-bodies, contain disorder-promoting amino acids and encode nuclear localised proteins. Finally, we identify ribosome pause sites that are resolved or induced by the depletion of CNOT1.

**Conclusion:** We define the key mRNA features that determine how the human Ccr4-Not complex differentially regulates mRNA fate and protein synthesis through a mechanism linked to codon composition, amino acid usage, and mRNA localisation.

## Introduction

The Ccr4-Not complex is a large evolutionarily conserved multi-protein complex (1–5), first described as a regulator of transcription (6–10). It has since been shown to have key regulatory roles extending well beyond transcription. It is a major complex involved in regulating an mRNA throughout the entire mRNA life cycle including: facilitating mRNA export (11), co-translational assembly of protein complexes (12), translational repression (13, 14), deadenylation and mRNA destabilisation (15–17). In humans, the Ccr4-Not complex has a molecular weight of ∼1.2MDa (18). The details of how these different regulatory outputs are exerted and whether they are target-specific is not fully understood. At the heart of the Ccr4-Not complex is the CNOT1 subunit, one of the roles of this subunit is to function as a scaffold to bring together the complex subunits as well as many additional effector proteins contributing to the diverse functions of the complex and thus acts as the central node for all the complex’s functions (16,18–22).

The Ccr4-Not complex has been shown to be delivered to mRNAs by multiple different mechanisms including: interaction with RNA-binding proteins that bind at the 3’UTR (23–26), recruitment to miRNA-bound mRNAs by the miRISC complex (27) and interaction directly of the Not5 subunit with the E-site of ribosomes with no tRNA present at the A-site (28). The most studied role of the Ccr4-Not complex is its involvement in mRNA deadenylation - the removal of the poly(A) at the 3’ end of the mRNA (29–32), which requires the activity of deadenylase subunits CNOT6/CNOT6L & CNOT7/CNOT8 (known as Ccr4 and Caf1, respectively, in yeast) (16, 33). This is the primary event in the mRNA decay pathway followed by the removal of the 5’ cap and subsequent degradation of the target mRNA (27,34–37). Deadenylation requires the expulsion of the poly(A) binding protein (PABP), which is thought to play an important role in the stability and translation of the mRNA (38).

The involvement of Ccr4-Not complex in the regulation of translation can be independent of its deadenylase activities (2). Indeed, deadenylation is not essential for translation repression of an mRNA by the Ccr4-Not complex in conjunction either with RBPs (39, 40) or miRNAs (41–44). Components of the Ccr4-Not complex are present in polysomes and are thought to be involved in translational quality control in yeast (45, 46). In addition, the complex has been implicated as a player in the buffering of gene expression (mechanisms that allow for compensatory regulation of mRNA levels and translation in the maintenance of protein homeostasis) (47, 48).

The role of the open reading frame and its sequence composition has recently emerged as equally important in the control of gene expression as that of the 5’ and 3’UTRs, which have traditionally been seen as the regulatory hubs of the mRNA. More specifically, codon usage has been highlighted as a key attribute linking translation elongation to mRNA stability (49–53). Interestingly, it is the nucleotide at the third “wobble” position of the codon that can confer stabilising / destabilising effects on the mRNA (54). The precise mechanisms linking codon usage to mRNA abundance, translation elongation and protein output are still not fully understood.

It is clear that the Ccr4-Not complex is a master regulator of mRNA fate. Despite this, no effort has been made to uncouple the impact of the Ccr4-Not complex on translation and mRNA abundance in shaping the final proteome on a system-wide level. Also, there has not been a global investigation of the mRNA features that predispose an mRNA to specific fate outcomes regulated by the Ccr4-Not complex. Here we employed a number of high throughput approaches - ribosome profiling, total RNA-seq, mRNA half-life studies and pulsed SILAC - in the context of depletion of the scaffold protein CNOT1 to understand the complex’s activity in post-transcriptional control. Knockdown of CNOT1 also downregulates the synthesis of many other subunits of the Ccr4-Not complex (19), which our results confirm (Additional File 1: Fig. S1A). We identify the features of mRNAs that determine the mechanism by which the Ccr4-Not complex regulates mRNA fate. Specifically, we uncover the importance of the precise codon composition of an mRNA in determining how the Ccr4-Not complex controls mRNA stability. Moreover, we uncover that mRNA localisation influences how the Ccr4-Not complex impacts mRNA translation: mRNAs translated at the ER are translationally downregulated after CNOT1 depletion, mRNAs that localise to p-bodies are translationally upregulated and mRNAs encoding proteins that localise to the nucleus are regulated at the level of translation but not stability by the Ccr4-Not complex. Lastly, we observe a role for the Ccr4-Not complex in the regulation of ribosome pause sites.

## Results

### Global increase of mRNA stability following CNOT1 knockdown

The Ccr4-Not complex is believed to be a major regulator of mRNA stability through its capacity to initiate mRNA decay by the deadenylase subunits, CNOT6/CNOT6L & CNOT7/CNOT8 (15,17,37,55,56). Recruitment of the Ccr4-Not complex is thought to occur on all mRNAs at the end of the transcript lifecycle. It has been shown for specific mRNAs that decay can be accelerated by the presence of particular RNA motifs that are bound by RBPs which then interact with Ccr4-Not complex subunits (57–62). However, this has not been tested on a global scale. To fully understand the distinct activities of the human Ccr4-Not complex in mRNA stability and translation, it was necessary to dissociate its roles in transcription from mRNA stability (2, 63), as both of these processes contribute to steady state mRNA levels (64).

To determine how the Ccr4-Not complex impacts mRNA half-lives transcriptome-wide, we sampled RNA at multiple time points after transcriptional inhibition using triptolide (65) with and without CNOT1 depletion. Titration experiments were used to determine the optimal concentration of triptolide (1μM), which produces a good decay curve for the short-lived MYC transcript without adversely affecting rRNA synthesis or cell viability (Additional File 1: Fig. S1B-D). RNA was isolated at 0, 0.5, 1, 2, 4, 8 and 16 hrs after transcriptional inhibition and the CNOT1 knockdown (Additional File 1: Fig. S1E) and RNA integrity (Additional File 1: Fig. S1F) were verified for each time point before performing RNA-seq.

Examination of RNA levels at each time point relative to the 0hr time point shows a global increase in mRNA stability following depletion of CNOT1 (Fig. 1A), clearly demonstrating the complex’s central role in mRNA destabilisation in human cells. An exponential model of decay y ∼ y_0_ e^-kt^ was fitted to the data, where k is the decay rate, y_0_ is the mRNA level at time point 0, and y_t_ is the mRNA level at time t. The mRNA half-life was then calculated using the equation: t_1/2_ = ln(2)/k. The half-lives obtained in control conditions for HEK293 cells showed a good correlation (r=0.406) with published half-lives from HEK293 cells obtained using a 4-thiouridine-based methodology (66). The median half-life in control conditions was 6.7 hrs and there was an average log2FC in half-life of 2.1 following CNOT1 depletion (Supplemental Table 1), demonstrating there is a substantial global increase in mRNA stability. Using k-means clustering the mRNAs can be grouped into three major clusters that are distinguishable by their half-lives in the presence of CNOT1 and the extent of mRNA stabilisation following CNOT1 depletion (Fig. 1BC). This demonstrates that the vast majority of mRNAs rely on the Ccr4-Not complex for mRNA turnover and there are some mRNAs which are particularly susceptible to rapid destabilisation by the Ccr4-Not complex. The mRNA half-lives for specific mRNAs from each cluster (Additional File 1: Fig. S2ABC) have been validated by qPCR with a different transcriptional inhibitor (flavopiridol) and a different pool of siRNAs targeting CNOT1 (Additional File 1: Fig. S2DEF). Overall, this shows that the Ccr4-Not complex is also the major regulator of mRNA stability in human cells and for the first time quantifies this on a global scale.

**Fig. 1:**
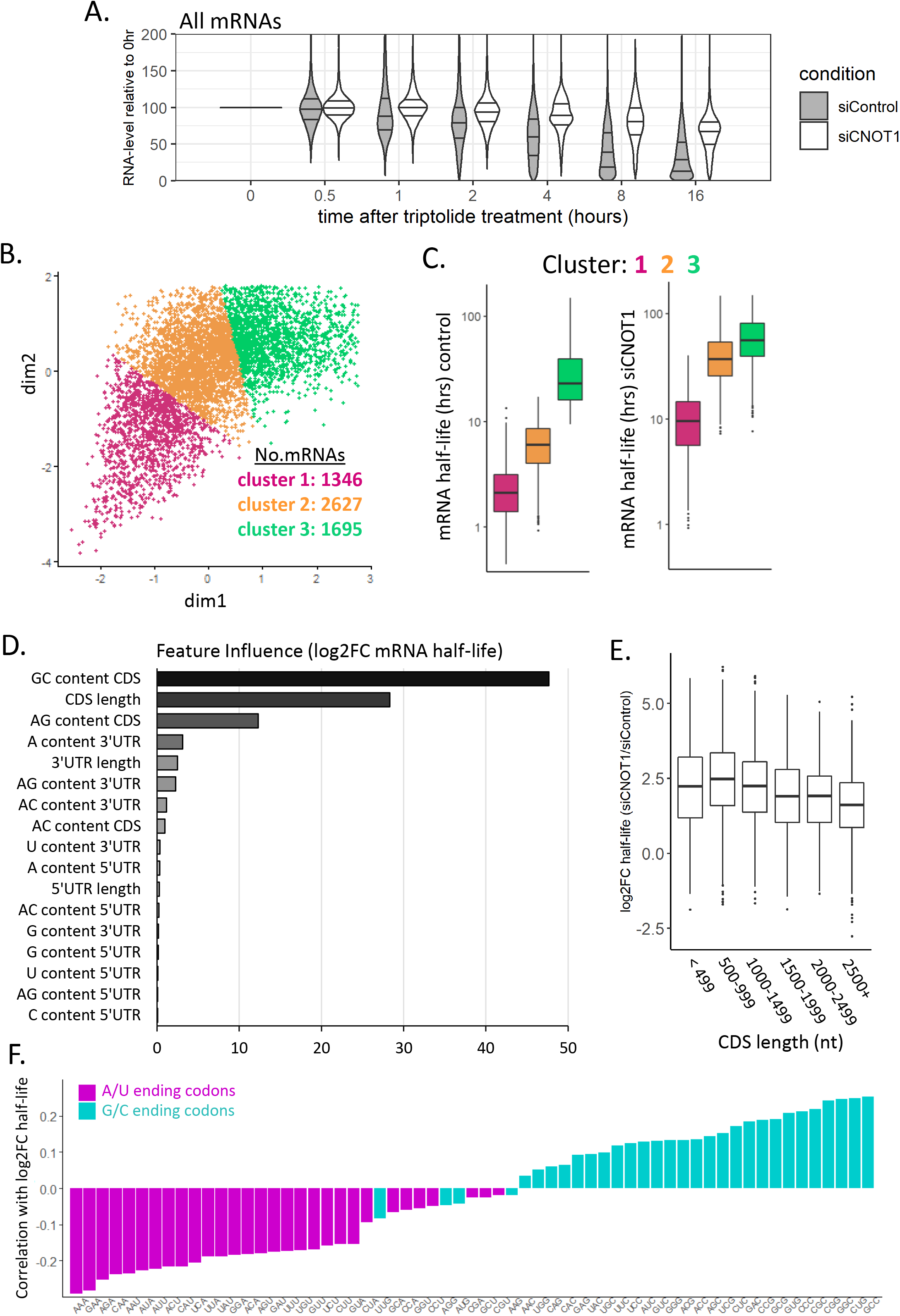
G/C-ending codons drive mRNA destabilisation by the Ccr4-Not complex. **A.** Plot of RNA abundance across multiple time points after inhibition of transcription with triptolide relative to the 0hr timepoint for siControl and siCNOT1 treated samples (three biological replicates). **B.** mRNAs grouped based on their mRNA half-life in the presence and absence of CNOT1 using k-means clustering. **C.** The mRNA half-lives before and after CNOT1 depletion for the three major clusters of mRNAs determined in (B). **D.** The variable influence (determined using gradient boosting) of mRNA sequence features on the log2FC mRNA half-life after CNOT1 knockdown. **E.** Log2FC mRNA half-life following CNOT1 depletion for mRNAs grouped by their CDS length. **F.** Correlation coefficient (Spearman’s Rho) between the frequency of a given codon in an mRNA and the log2FC mRNA half-life (siCNOT1 / siControl). Codons with an A/U at the 3^rd^ nucleotide position are coloured in magenta and codons with a G/C at the 3^rd^ nucleotide position are coloured in cyan.

Moreover, gene ontology analysis conducted on all of the mRNAs ranked by their log2FC half-life after CNOT1 knockdown showed a significant enrichment for only a small number of terms, likely due to the fact that the Ccr4-Not complex is the major regulator of the stability of most mRNAs. The terms that were significant related to mitochondrion organisation, cardiac/muscle development and regulation of S/T kinase activity (Additional File 1: Fig. S3A). These observation may point towards a mechanism behind recent findings that the depletion of CNOT1 delays neurodevelopment (67) and affects cardiac function (68), in that, these specific subsets of mRNAs are heavily reliant on the Ccr4-Not complex for their stability.

### G/C-ending codons drive Ccr4-Not mediated mRNA destabilisation

There are a number of attributes of the mRNA sequence that impact mRNA translation or stability including: length, mRNA structure and nucleotide composition within different regions of the mRNA (69, 70). To determine if any of these are potentially involved in directing Ccr4-Not regulation of mRNA stability, the contribution of these variables to the change in mRNA half-life following CNOT1 knockdown was evaluated using gradient boosting (71). First mRNA features were pre-filtered to remove highly correlated variables (r>0.7, Additional File 1: Fig. S3B). Analysis of the independent variables showed that GC content and the length of the CDS have the greatest influence on the change in mRNA half-life (Fig.1D). Grouping of mRNAs by CDS length shows that shorter CDSes are associated with greater mRNA stabilisation following CNOT1 knockdown (Fig. 1E).

The defining aspect of CDS sequence composition is that it is organised into codon triplets. CDS GC content is highly correlated with the GC content of the 3^rd^ nucleotide of the codon (Additional File 1: Fig. S3C) and components of the Ccr4-Not complex have previously been implicated in codon-mediated regulation of mRNA translation in yeast and zebrafish (16,72,73). Therefore, we hypothesised that the observed differences in human cells may be driven by the codon usage within the mRNAs.

The codon stabilisation coefficient is the correlation between the codon frequency and the half-life of an mRNA (74). Here we determined the correlation of the frequency of a given codon with the log2FC in mRNA half-life after CNOT1 depletion. Strikingly, this shows a strong split between the correlations of codon frequency of G/C-ending (cyan) and A/U-ending (magenta) codons with the change in mRNA half-life (Fig. 1F). In general, the greater the frequency of any given G/C-ending codon, the greater the increase in half-life after CNOT1 knockdown (Fig. 1F). Together, this supports the presence of G/C-ending codons is a primary driver of destabilisation of an mRNA via the Ccr4-Not complex.

Synonymous codons are those which differ in sequence but encode the same amino acid. The distinct transcript pools present in proliferation and differentiation have been shown to have opposing synonymous codon usage signatures (75–77). mRNAs enriched for A/U-ending codons are abundant in proliferation, whereas it is the mRNAs that contain more G/C-ending codons that are abundant in differentiation (76, 77). Having observed a clear distinction in how G/C-ending and A/U-ending codons impact Ccr4-Not-mediated regulation of an mRNA’s stability (Fig. 1F), we sought to understand if this was driven by synonymous codon usage differences. Hence, the correlation of log2FC half-life with codon frequency (Fig. 1F) was reordered by the amino acid (Additional File 1: Fig.S3D). Unexpectedly, this highlights that while synonymous codons are important for the distinction of mRNA stability regulation by the Ccr4-Not complex, the amino acid itself further impacts the change in mRNA half-life with CNOT1 knockdown (Additional File 1: Fig. S3D). For example, the G/C-ending codons of some amino acids (e.g. Ala & Pro) correlate with an increase in mRNA stability, but their synonymous A/U-ending codons show very minimal correlation with stability changes (Additional File 1: Fig. S3D). A recent publication highlighted that amino acid composition also affects mRNA stability (78) and here we expand on this showing how amino acid differences contribute to the nature of the impact of G/C or A/U ending codons on mRNA stability regulation by the Ccr4-Not complex.

### Translational regulation by the Ccr4-Not complex

It is proposed that the regulation of mRNA stability is linked to translation elongation (16,52–54,79,80). Recent structural data from yeast shows the Not5 subunit of the Ccr4-Not complex can interact directly with the ribosome, this interaction occurs at the ribosomal E-site when the A-site is unoccupied (28). In addition to its described role in mRNA destabilisation, the Ccr4-Not complex has also been implicated in the regulation of translational repression, which can occur independent of deadenylation (39,41,81–83). As of yet no study has investigated the role of the human Ccr4-Not complex at the level of translation on a system-wide level. Polysome gradients show there is a global accumulation of polysomes following CNOT1 depletion (Fig. 2AB). Whether the translational upregulation following CNOT1 depletion is a direct consequence of the global increase in mRNA stability (Fig. 1A) or if the role of the Ccr4-Not complex in the regulation of translation is distinct from how it controls mRNA stability is unknown. Therefore, to assess the impact of the Ccr4-Not complex on translation of individual mRNAs globally at codon resolution, ribosome profiling was conducted with and without the depletion of CNOT1 (Fig. 2AB, Additional File 1: Fig.S4A-C). Ribosome profiling involves high throughput sequencing of ribosome protected fragments (RPFs) (84, 85). Quality control analysis of the RPF sequencing data showed the three replicates contained an extremely low number of reads aligning to rRNA (Additional File 1: Fig. S5A), were highly correlated (Additional File 1: Fig. S5B), had the expected read length distribution (Additional File 1: Fig. S5C), the majority of reads aligned to the CDS (Additional File 1: Fig. S5D) and showed a strong periodicity (Additional File 1: Fig. S5E). This demonstrates the ribosome profiling data is of very high quality and provides a benchmark ribosome profiling dataset.

Translational efficiency (TE) is defined as the number of RPFs aligning to a given CDS corrected for the mRNA’s abundance (determined by parallel total RNA-seq). Due to differences in library preparation for RPFs and total RNA, differential expression analysis using DESeq2 (86, 87) was conducted separately for each library type to obtain a log2 fold change for RPF and RNA changes independently (Supplemental Table 2). The change in translational efficiency was then determined by calculating log2FC RPF – log2FC RNA. Our mRNA stability experiments demonstrated the global upregulation of mRNA stability after CNOT1 depletion (Fig. 1A). There is also a global increase in mRNA ribosome occupancy following CNOT1 depletion (Fig. 2B). Therefore, in this condition, and due to the nature of differential expression analysis, it is more appropriate in this slightly unusual context to consider a negative log2FC as least upregulated and the positive log2FC as most upregulated. Using the ribosome profiling, three translationally regulated groups were defined (Fig. 2C): mRNAs with an increase in RPFs complementary to the increased mRNA stability thus having no effective TE change (yellow); mRNAs with a translation increase greater than the mRNA stability increase (increased TE; red) and mRNAs with a translational increase lower than the mRNA stability increase (decreased TE; blue). To validate the translational observations for individual mRNAs within these groups (Fig. 2DEF & Additional File 1: Fig. S6), independent experiments (n=2) were conducted without cycloheximide pre-treatment, and RT-qPCR applied across all fractions of polysome gradients following treatment with CNOT1 or control siRNA. Using this alternative approach, we show that mRNAs with altered translational efficiency display the expected changes in polysome distribution (Fig. 2DF, Additional File 1: Fig S6ABEF). In addition, mRNAs with correlated changes at both the RPF level and RNA level (no effective TE change, yellow) show no major changes in distribution across polysomes (Fig. 2E, Additional File 1: Fig. S6CD). Together this confirms the ribosome profiling analysis correctly identifies mRNAs with altered translation after CNOT1 knockdown.

**Fig. 2:**
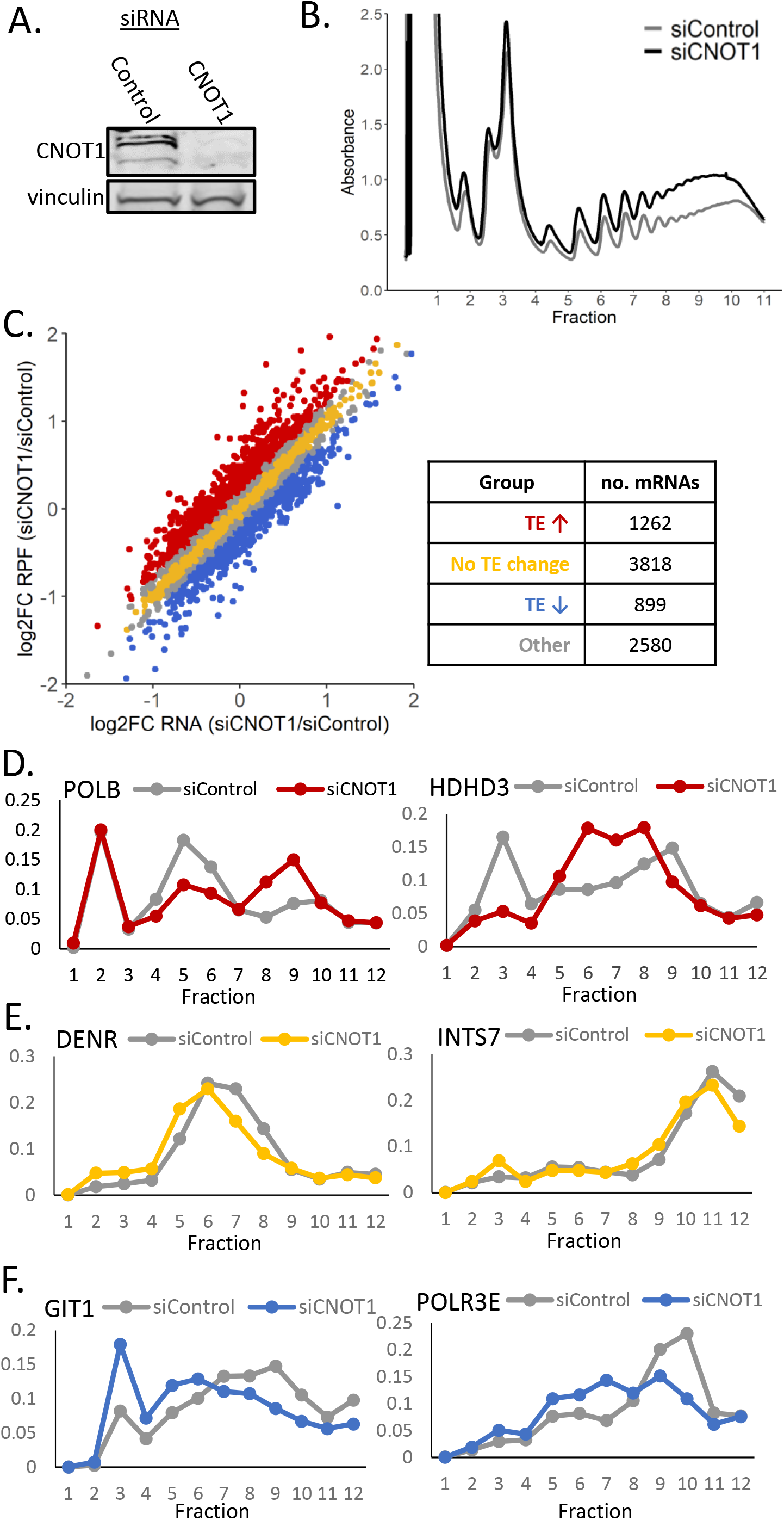
Ribosome profiling identifies mRNAs regulated by the Ccr4-Not complex at the level of translation. **A.** Western blot confirms siRNA knockdown of CNOT1. Vinculin is used as a loading control. **B.** Polysome gradient profiles for samples treated with control or CNOT1-targeting siRNA. **C.** There are groups of mRNAs with distinct changes in ribosome occupancy and mRNA abundance when CNOT1 is depleted. Log2 fold change of RPFs and RNA following CNOT1 depletion were determined using DESeq2 independently for each library type. Log2 translational efficiency (TE) was determined by log2FC RPF – log2FC RNA; a threshold of log2FC TE > 0.2 was used to categorise mRNAs as having increased TE and a log2FC TE < -0.2 for mRNAs with decreased TE. No TE change was classified by a log2TE <0.1 & >-0.1. The table shows the number of mRNAs present in each group. **DEF.** qPCR along gradient fractions from an independent experiment with and without CNOT1 depletion (n=1 shown, n=2 is shown in Additional File 1: Fig. S6). **D.** Validation of the increased TE after CNOT1 depletion of POLB and HDHD3 from the group of mRNAs identified in (C, red). In grey is the control and in red is the CNOT1 siRNA treated. **E.** Validation of the unchanged TE after CNOT1 depletion for DENR and INTS7 from the group of mRNAs identified in (C, yellow). In grey is the control and in yellow is the CNOT1 siRNA treated. **F.** Validation of the decreased TE after CNOT1 depletion of GIT1 and POLR3E from the group of mRNAs identified in (C, blue). In grey is the control and in blue is the CNOT1 siRNA treated.

### Differentially translated mRNAs are functionally distinct

To determine if the functions of the proteins encoded by the mRNAs regulated at the level of translation by the Ccr4-Not complex are different, gene set enrichment analysis was conducted for the mRNAs ranked by the extent of the TE change with CNOT1 depletion (Fig. 3A, Additional File 1: Fig. S7AB, Supplemental Table 3). This showed a large number of gene ontology (GO) terms associated with decreased TE after CNOT1 knockdown. This included GO terms related to development & morphogenesis, cell signalling pathways and structural components of the cell (Additional File 1: Fig. S7A). These mRNAs are also associated with the endoplasmic reticulum (ER), extracellular matrix (ECM) and plasma membrane (Fig. 3A). To investigate this in more detail, cell lysates were separated into cytosolic and ER fractions (Fig. 3B) and the RNA present in each fraction sequenced. K-means clustering has been used to define mRNAs that predominantly localised to the cytosol or ER (Fig. 3C, Supplemental Table 4); using these groups shows a greater proportion of mRNAs translationally downregulated following CNOT1 depletion are indeed ER-targeted mRNAs in HEK293 cells (Fig. 3D) and the proteins encoded are localised at the ER and plasma membrane (Additional File 1: Fig.S7CD, data from U2OS cells (88)).

**Figure 3:**
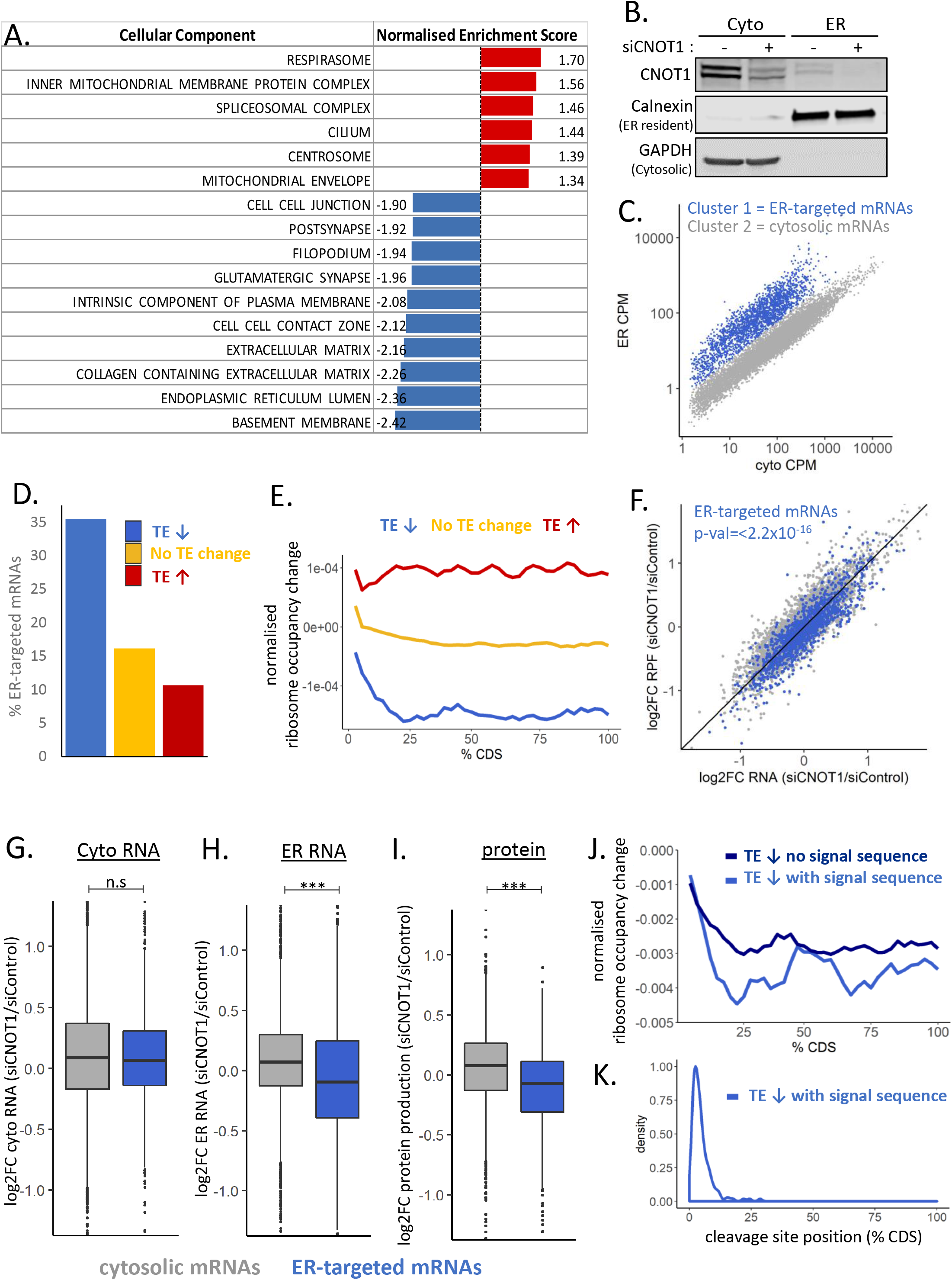
ER-targeted mRNAs are translationally downregulated following CNOT1 depletion. **A.** All mRNAs were ordered by the log2FC TE (siCNOT1/siControl) and gene set enrichment analysis conducted on the ranked list using the fgsea R package for cellular components. **B.** Western blot confirming lysate fractionation into ER and cytosol with and without the depletion of CNOT1. **C.** k-means clustering of mRNAs based on their abundance (CPM) in the ER and cytosolic fractions defines two groups of mRNAs based on their predominant localisation. **D.** mRNAs with decreased TE when CNOT1 is depleted (identified in Fig. 2C) are enriched for targeting to the ER in control conditions. **E.** Median change in ribosome occupancy (normalised for mRNA abundance) assessed across the CDS for the groups of mRNAs identified in Fig. 2C. **F.** ER-targeted mRNAs globally decrease translationally efficiency following CNOT1 depletion.. **G.** log2FC RNA in the cytosolic fraction between CNOT1 depleted and control conditions. **H.** log2FC RNA in the ER fraction between CNOT1 depleted and control conditions shows a reduction in mRNAs that are targeted to the ER in control conditions localising to the ER after CNOT1 depletion. **I.** Displayed is the protein production change (siCNOT1/siControl) for mRNAs predominantly localised to the cytosol or the ER. For FGHI significance was determined using the Kruskal-Wallis test. **J.** Median change in ribosome occupancy (normalised for mRNA abundance) assessed across the CDS for the mRNAs with decreased TE after CNOT1 depletion separated into two groups based on the predicted presence of a signal sequence (determined using SignalP (92)). 157/899 mRNAs were predicted to contain a signal sequence. **K.** Predicted location of the signal peptidase cleavage site in the group of mRNAs with a predicted signal sequence (determined using SignalP (92)).

In contrast, mRNAs encoding proteins associated with the mitochondria, splicing or the centrosome are translationally upregulated following CNOT1 depletion (Fig. 3A, Additional File 1: Fig. S7A). Increased TE is also associated with proteins involved in tRNA processing and modification as well as proteins having molecular functions involved in DNA binding and repression of transcription (Additional File1: Fig.S7AB). Overall, this analysis suggests a role for the Ccr4-Not complex in the translational regulation of functionally distinct and spatially localised groups of mRNAs.

### Decreased ribosome occupancy after CNOT1 depletion occurs downstream of signal sequences

Next, we examined the change in ribosome occupancy across the CDS. This showed for mRNAs with increased translational efficiency the increased ribosome occupancy is evenly distributed across the CDS (Fig. 3E). However, for translationally downregulated mRNAs ribosome occupancy ramps down sharply from the start codon throughout the first ∼10% of the CDS, followed by a large and even reduction across the final 75% of the CDS (Fig. 3E). This suggests these mRNAs require the presence of the Ccr4-Not complex for their efficient translation in the first section of their CDS in control conditions. For example, this may be the result of the presence of regulatory sequences in this region that control mRNA localisation, such as the signal sequence recognised by the signal recognition particle (SRP), which would conform with our observation about the high abundance of ER-targeted mRNAs in this group (Fig. 3D). In addition, we see a highly significant global reduction in the translational efficiency of ER-target mRNAs following CNOT1 depletion (Fig. 3F), suggesting the Ccr4-Not complex specifically plays a role in the regulation of mRNAs that localise to the ER to be translated.

To investigate this further we fractionated cells into the ER and cytosol with and without CNOT1 depletion (Fig. 3B) and sequenced the RNA from each fraction. Using the classification of mRNAs predominantly localised in the ER or cytosol in control conditions (Fig. 3C, Supplemental Table 4) we were able to assess how these mRNAs change localisation after CNOT1 knockdown. This clearly shows that ER-targeted mRNAs have reduced levels specifically in the ER after CNOT1 knockdown (Fig. 3GH).

To confirm the impact of the altered mRNA localisation and translational efficiency of ER-targeted mRNAs on protein output, pulsed SILAC (stable isotope labelling by amino acids in culture (89)) was conducted following CNOT1 depletion (Additional File 1: Fig. S8A). A protein was only included in the analysis if it was detected in both the forward and reverse labelling technical replicates (Additional File 1: Fig. S8) and in at least two of the three biological repeats, this resulted in a group of 3495 proteins (Additional File 1: Fig. S8C, Supplemental Table 5). The pulsed SILAC confirms that the reduced TE of ER-target mRNAs is reflected in reduced protein synthesis (Fig. 3I).

Ribosome pausing can occur at the signal sequence (90) and if the mRNA is not correctly translocated to the ER for the continuation of its translation, this would result in lower ribosome occupation of the latter part of the CDS – as we observe. A very recent publication observed disome populations at these signal sequences (91), and our data suggests the involvement of the Ccr4-Not complex in the regulation of these pause sites. To investigate this in more detail SignalP (92) was used to select mRNAs with predicted signal sequences. Separation of the positional data by the presence or absence of predicted signal sequence shows sharper decline in ribosome occupancy for mRNAs with a predicted signal sequence (Fig. 3J). The SRP cleavage site, on average, is positioned at around 4.75% of the length of the CDS (Fig. 3K). This position coincides with the sharp decline in ribosome occupancy in the absence of CNOT1. This might suggest that the Ccr4-Not complex accelerates the progression of the ribosome through this site by either facilitating localisation of the mRNA or the efficiency of cleavage. Thus, in the absence of the complex, ribosome occupancy downstream of this position is significantly diminished because the ribosomes cannot progress.

### The impact of codons on Ccr4-Not complex mediated regulation of translational efficiency

Having observed that an increase in mRNA half-life after CNOT1 depletion positively correlates with the frequency of G/C-ending codons (Fig.1F), the correlation of codon frequencies with the log2FC TE was next examined. This showed it is A/U-ending codons that positively correlated with an increase in TE after CNOT1 knockdown (Additional File 1: Fig. S9A). However, of note is the extent of these correlations which is not as strong as observed for mRNA half-life (Fig.1F). Nevertheless, there is a strong distinction in the direction of the correlation based on the 3^rd^ nt of the codon as indicated in magenta/cyan (Additional File 1: Fig S9A). To confirm the influence of this factor the luciferase reporter system was utilised. The Renilla luciferase CDS is naturally rich in A/U-ending codons (74.5%), hence the three G/C-ending codons (AUC, GUC & ACC) most negatively correlated with the TE change were substituted into the Renilla sequence at the corresponding synonymous codon positions. The translational efficiency was then determined using the luciferase activity / luciferase RNA level determined by qPCR with firefly luciferase used as a transfection control. This reporter clearly demonstrates that conversion of A/U-ending codons in the Renilla CDS to synonymous G/C-ending codons leads to a reduction in the extent of TE change after CNOT1 knockdown (Additional File 1: Fig. S9B).

mRNAs with AU-rich CDSes and 3’UTRs have been shown to be enriched in p-bodies (sites of translational repression and mRNA storage) (93) and CNOT1 depletion prevents p-body formation (19). We are able to clearly show, using the previously published HEK293 p-body transcriptome (94), that mRNAs translationally upregulated after CNOT1 knockdown are most enriched in p-bodies in control conditions (Additional File 1: Fig. S9C). The exact nature of p-bodies is not fully understood; they contain components of the deadenylation and decapping machinery (95–101) and the role of deadenylation in their formation is debated (57,93,94,102–106). Our finding suggests that these specific mRNAs might undergo targeted translational repression by the Ccr4-Not complex followed by subsequent shuttling to p-bodies for storage.

### How regulation of mRNA translation and/or stability by the Ccr4-Not complex impacts protein output

As we had determined that there is a global upregulation of both mRNA stability and translation following depletion of CNOT1, we next sought to understand the role of the Ccr4-Not complex in translation both alongside and independent of its role regulating mRNA stability. To dissect how the Ccr4-Not complex regulates mRNA translation compared to its control of stability, mRNAs were grouped based on the change in their translational efficiency (as assessed by ribosome profiling) and change in mRNA half-life (determined by triptolide inhibition) following CNOT1 depletion (Fig. 4A). This generates six groups of mRNAs classified first by whether there is decreased TE, no TE change or increased TE (blue, yellow & red, respectively, as in Fig. 2C), and second by a small change in mRNA half-life or a large increase in mRNA half-life (dark, light, respectively) when CNOT1 is depleted (Fig. 4A; Additional File 1: Fig. S10A).

**Fig. 4:**
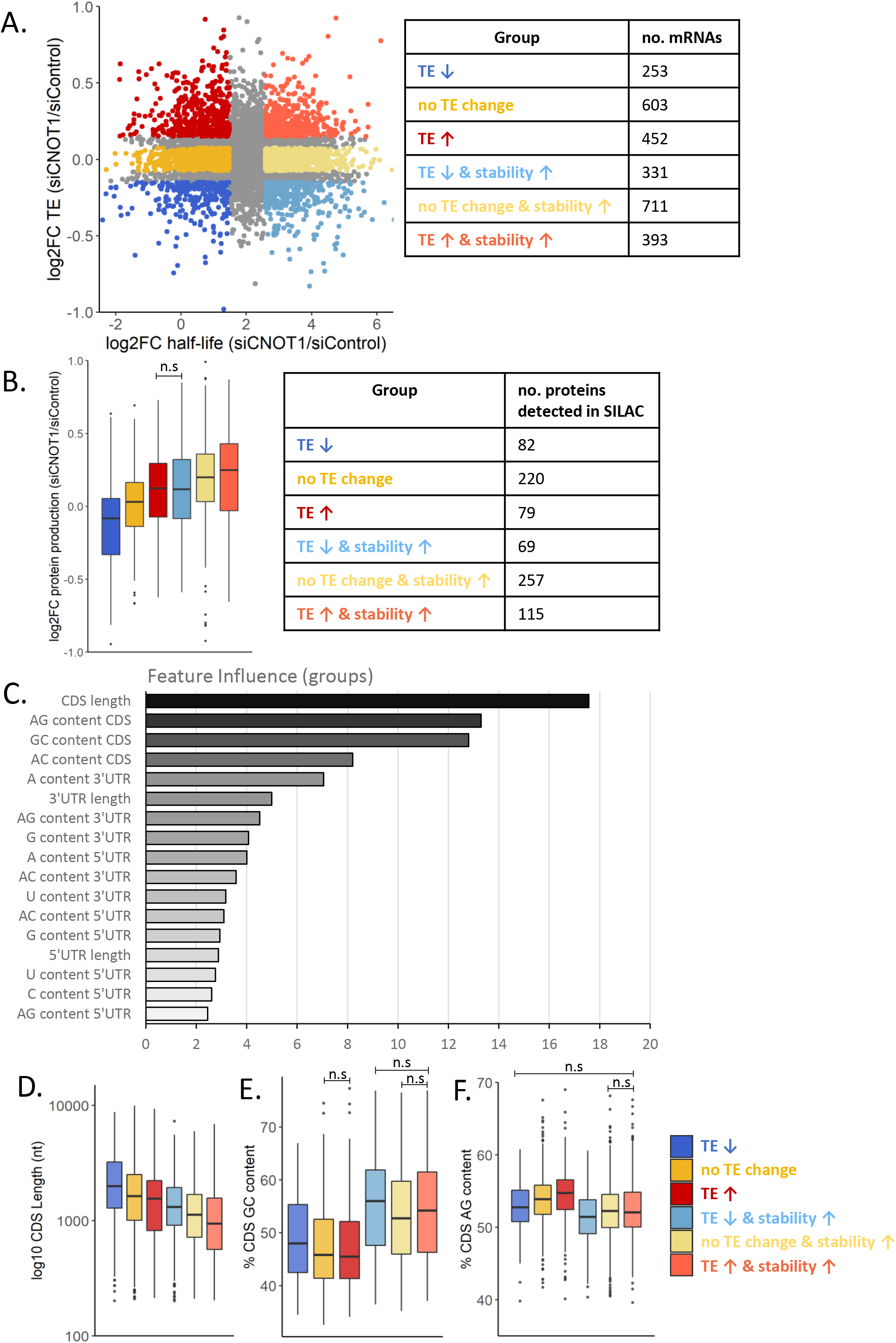
mRNA features that distinguish the role of the Ccr4-Not complex in the regulation of mRNA translation v stability. **A.** Data shows the log2FC in mRNA half-life after CNOT1 depletion (x-axis) compared to the log2FC translational efficiency (y-axis). mRNAs are first classified based on their mRNA stability change – a low log2FC mRNA half-life (dark colours) or a high log2FC mRNA half-life (light colours) and second by their TE change – increased TE (red), no TE change (yellow) or decreased TE (blue). **B.** Pulsed SILAC data for three biological repeats conducted with forward and reverse labelling (Additional File 1: Fig. S8A-C). Displayed is the protein production change (siCNOT1/siControl) for the proteins that were detected in at least two biological repeats for the groups of mRNAs classified in A. Non-significant comparisons are indicated, and all statistical comparisons are shown in Additional File 2: Fig. S1A. **C.** The influence of mRNA sequence features on the classification of mRNAs into groups based how the Ccr4-Not complex differentially regulates their stability and/or translation (Fig. 4A) was determined by gradient boosting. **DEF.** The presence of the top three features identified in **D** in the differentially regulated groups of mRNAs. **D.** CDS length, **E.** CDS G/C nucleotide content & **F.** CDS A/G nucleotide content. Statistical comparisons for BCD are shown in Additional File 2: Fig. S1B-D. For clarity non-significant comparisons are indicated, and all statistical comparisons are shown in Additional File 2: Fig. S1BCD.

Next the aim was to investigate how the observed translation and stability changes impact protein output by use of the pulsed SILAC data (Additional File 1: Fig. S8). The Ccr4-Not complex has a global role in the regulation of both mRNA stability and translation, and this is the first time the influence of this complex on protein output has been assessed. Analysis of protein level changes in the differentially regulated groups of mRNAs (Fig.4A). This shows that increased stability after CNOT1 knockdown results in increased protein synthesis compared to mRNAs with a minimal change in stability (light v dark colours, Fig. 4B). In addition, increased TE after CNOT1 knockdown is associated with increased protein synthesis (red v yellow, Fig. 4B) and decreased TE associated with decreased protein synthesis in comparison to mRNAs with a similar mRNA stability change but no change in TE (blue v yellow, Fig. 4B).

### CDS composition differentiates how the Ccr4-Not complex regulates of mRNA translation vs stability

The Ccr4-Not complex has been shown to have roles in the regulation of both mRNA stability and translation (13–17,19,107). To identify mRNA features that specifically influence how the complex regulates translation as opposed to stability, the importance of these variables in determining the mRNA group assignment (classification as in Fig. 4A) was evaluated using gradient boosting (71). The feature analysis again points toward the CDS as a major driver for differential regulation of mRNA fate mediated via the Ccr4-Not complex, with the four most influential features pertaining to the CDS (Fig. 4C). Closer analysis of the top influential features shows it is shorter mRNAs that are most highly upregulated in terms of stability (Fig. 4D). The CDS GC content strongly distinguishes mRNAs with a large increase in half-life following CNOT1 depletion, from those with a lesser increase in half-life (Fig. 4E). Additionally, the AG content of the CDS distinguishes the translational changes between the groups of mRNAs with small increase in half-life (blue/yellow/red, Fig. 4F). mRNAs with increased translation are more AG-rich in the CDS than mRNAs with decreased translation (Fig. 4F).

In terms of 3’UTR features, the 3’UTR A content and length were the most influential for group classification (Fig. 4C). mRNAs with shorter 3’UTRs have increased TE with CNOT1 knockdown and mRNAs with decreased TE have longer 3’UTRs, with no influence of stability (Additional File 1: Fig. S10B). This suggests the translationally downregulated mRNAs may be more highly regulated as longer 3’UTRs means increased potential for the presence of regulatory sequences. Also, mRNAs with increased TE have a higher 3’UTR A content compared to the mRNAs with decreased TE that have a comparable half-life change (Additional File 1: Fig. S10C).

Overall, this analysis highlights a significant role for the CDS in coordinating Ccr4-Not complex function and further dissects the role of the complex in the regulation of translation compared to its roles in mRNA stability.

### Nuclear proteins enriched for disorder-promoting AAs are translationally upregulated

We have shown that the frequency of G/C-ending codons is associated with mRNA destabilisation by the Ccr4-Not complex (Fig. 1F) and feature analysis highlights the CDS composition as a distinguishing factor between differentially regulated groups of mRNAs (Fig. 4C-F). Analysis of average synonymous usage of codons for each of the groups of mRNAs revealed a strong distinction in the use of synonymous codons between mRNAs with a small and large increase in stability (Fig. 5A), in agreement with the earlier correlation data based on codon frequency (Fig 1F). However, there are no additional major differences in synonymous codon usage preferences when dissecting this further to the level of altered translation (Fig. 5A). This suggests the synonymous codon usage is involved in how the Ccr4-Not complex regulates mRNA stability but is not a major determinant of its independent role in translation. Further investigation of how codon frequencies ultimately impact protein synthesis demonstrates that although the 3^rd^ nucleotide preference of codons contributes to mRNA stability and translational efficiency regulated by the Ccr4-Not complex (Fig.1F & Additional File 1: Fig. S9A), their frequency does not strongly correlate with protein production changes after CNOT1 depletion (Additional File 1: Fig.S9D).

**Fig. 5:**
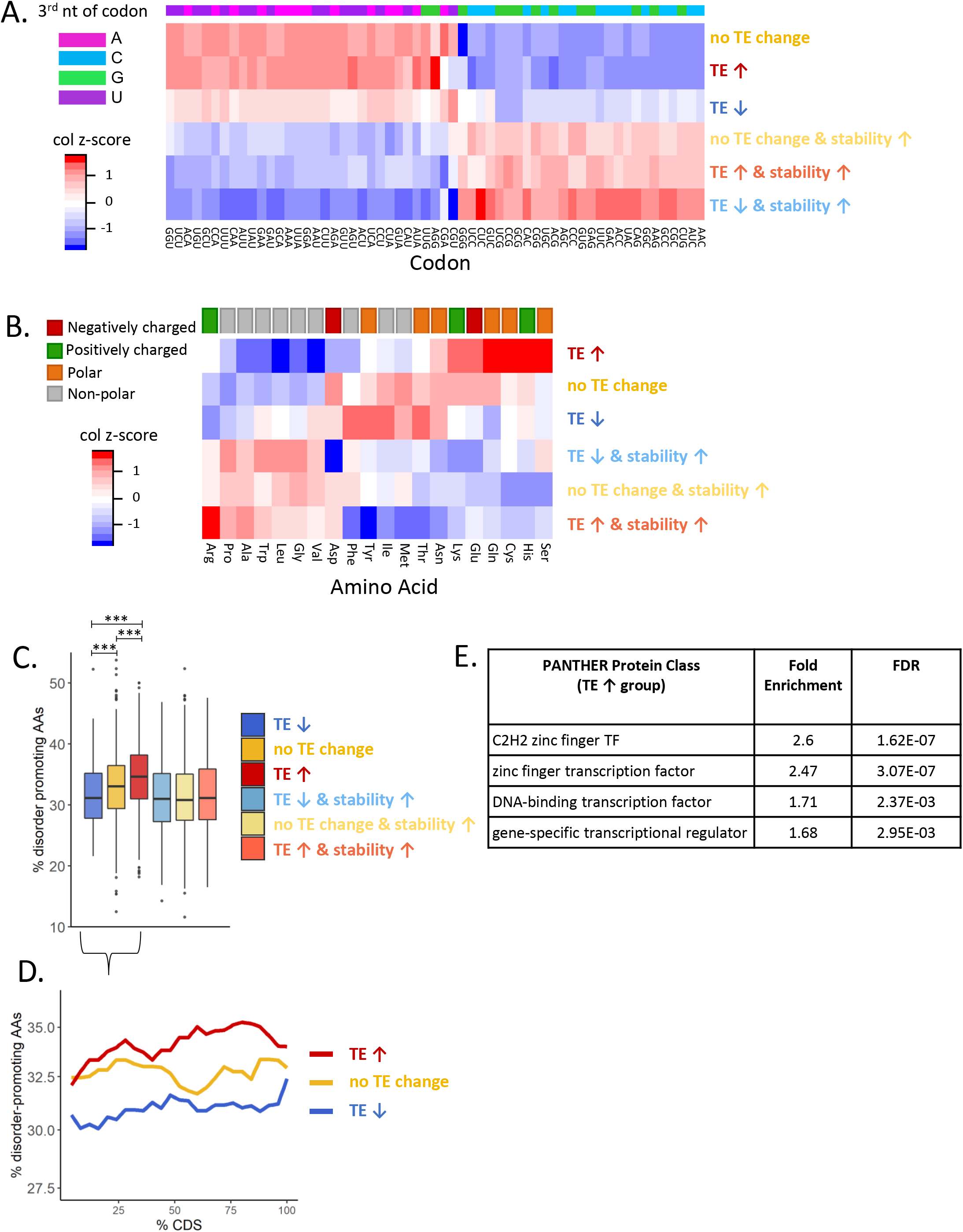
Nuclear proteins enriched for disorder-promoting AAs are translationally upregulated. **A.** Heatmap represents the synonymous codon usage of each codon that encodes a given amino acid and a column z-score has been applied to compare the codon preference between the groups of mRNAs. The coloured bar at the top indicates the nucleotide present at the third position in the codon. **B.** Heatmap shows the average frequency of each amino acid per mRNA and a column z-score has been applied. The coloured bar at the top classifies the amino acids by their side chain type. **C.** The percentage of amino acids encoded by the mRNAs that promote disorder in protein structure (108, 109). Key significant comparisons are shown, and all statistical comparisons are presented in Additional File 2: Fig. S2A. **D.** The localisation of disorder-promoting amino acids across the CDS for the groups of mRNAs with minimal changes in mRNA stability. **E.** Protein class enrichment analysis (using PANTHER (112)) shows that the mRNAs with increased TE but minimal increase in stability are enriched for transcription factors, specifically C2H2 zinc finger proteins.

We therefore examined the amino acid frequency in the differentially regulated groups of mRNAs. This shows, in addition to the synonymous codon usage, there is also a bias in the amino acid composition of these mRNAs (Fig. 5B). mRNAs with increased translation are enriched for polar/charged amino acids (AAs) and are depleted of non-polar amino acids (Fig. 5B). mRNAs with decreased translation show no preference for these same charged/polar amino acids (Fig. 5B). This suggests that it is a combination of codon and amino acid usage that influence how the Ccr4-Not complex regulates mRNA stability and translation and ultimately protein output.

Polar/charged AAs are often classified as disorder-promoting AAs in terms of their role in protein structure (108, 109). Therefore, we examined the presence of these disorder-promoting AAs in the differentially regulated groups of mRNAs, which clearly showed they are enriched in the mRNAs with increase TE but minimal change in stability (Fig. 5C). AAs are important for protein function and can influence ribosome decoding speeds (79,110,111). To understand the role of the disordered AAs in this group of mRNAs we assessed the localisation of the disordered AAs along the CDS. This showed the disordered AAs in the translationally upregulated group of mRNAs are enriched across the CDS and are particularly enriched at the 3’ end of the CDS compared to mRNAs with no effective TE change (Fig. 5D). Analysis of the protein class (112) of the mRNAs with increased TE showed an enrichment for transcription factors (TFs), specifically C2H2 zinc finger TFs (Fig. 5E) and protein localisation data confirms these translationally upregulated mRNAs encode proteins that are nuclear/chromatin localised (Additional File 1: Fig.S10D, data from (88)).

Unexpectedly, these zinc-finger protein mRNAs also have relatively short half-lives in control conditions, but do not show any change in mRNA stability with CNOT1 depletion (Additional File 1: Fig. S10E). This is interesting because the mRNA turnover of this group of mRNAs appears completely independent of the Ccr4-Not complex, suggesting they are regulated by a distinct decay pathway and the Ccr4-Not complex is only involved in their translational regulation, perhaps linked to their specialised role in transcriptional control.

### Ribosome pause sites regulated by the Ccr4-Not complex

The number of ribosomes on the mRNA at a point in time, has often been used as an indicator of protein output – the more ribosomes on the mRNA the more translated the mRNA and hence the more protein produced. However, this is not always the case, both elongation and initiation rates dictate the overall ribosome occupancy observed at a single point in time. For example, decreased ribosome occupancy could be an indicator of removal of an elongation block resulting in increased elongation speed and therefore fewer ribosomes on the mRNA. Conversely, increased ribosome occupancy could be a result of either resolving a block at initiation of translation or decreased elongation speed which would result in slower run-off of ribosomes.

In yeast it has been suggested that the Ccr4-Not complex is linked to ribosome pausing (12, 28). Our ribosome profiling data with and without depletion of CNOT1 in HEK293 cells was utilised to examine this further in human cells. First, a pause site in each condition was defined as a position with a RPF peak height ten times greater than the average RPF peak on the mRNA. Second, these pause sites were then classified as either ‘sustained’ meaning they are present but unaltered with CNOT1 knockdown; ‘resolved’ in that the reduction in RPF peak height is ten times greater than the average delta decrease across the mRNA or ‘induced’ in that the increase in RPF peak height is ten times greater than the average delta increase (Fig. 6A, Supplemental Table 6). Figures 6BCD show ribosome P-site occupancy for example mRNAs after normalisation for mRNA abundance in control and CNOT1 knockdown conditions that contain the distinct pause site types (ribosome P-site shown and determined by read offset of 12nt: Additional File 1: Fig. S11AB). It is possible for a specific mRNA to have pause sites meeting more than one of these criteria, Fig. 6A indicates the number of mRNAs containing combinations of these pause types and the mRNAs distinctly with one type are used in downstream analysis to characterise these pause site types in more detail.

**Fig. 6:**
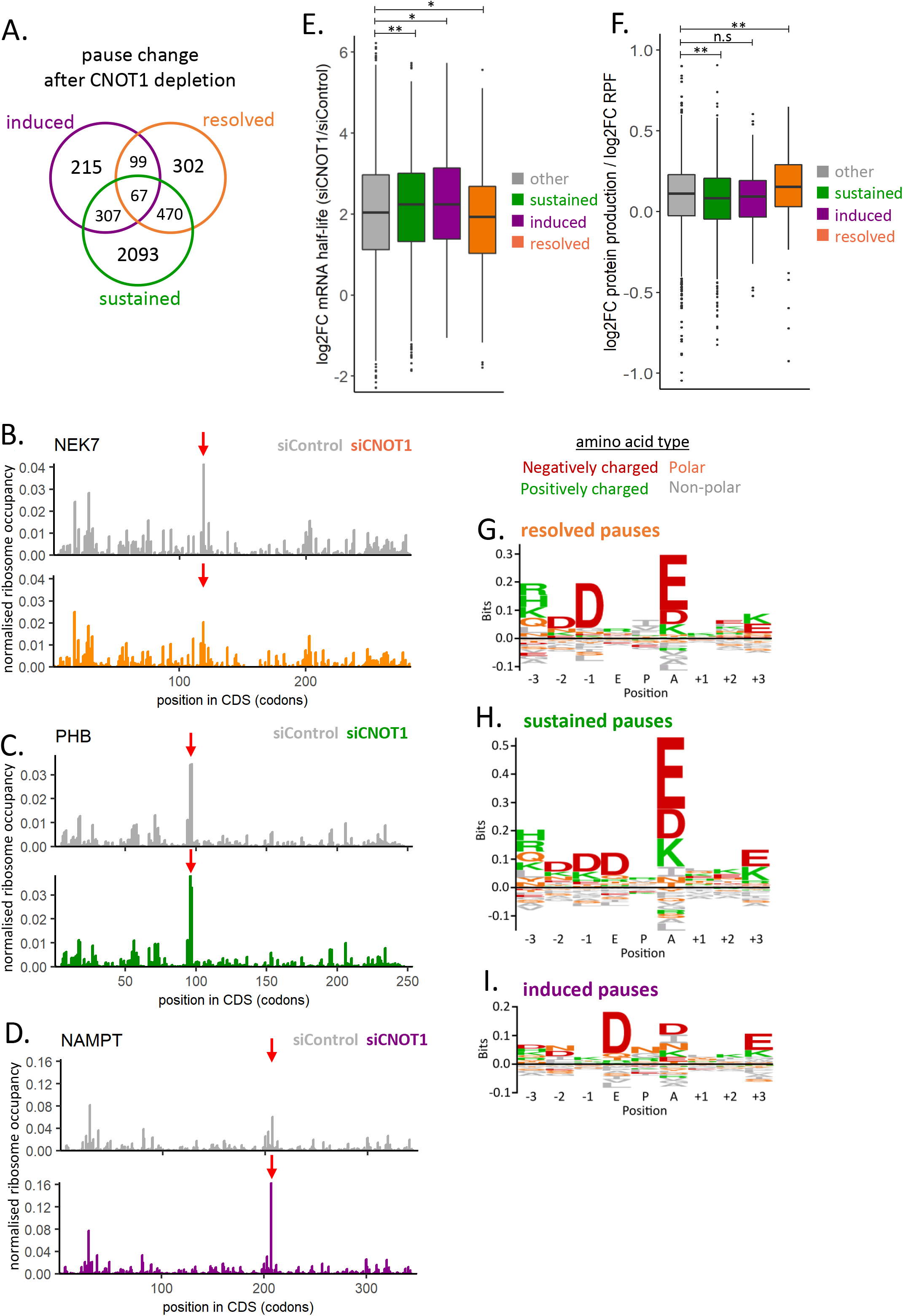
Ribosome pause sites regulated by the Ccr4-Not complex. **A.** Venn diagram indicates the number of mRNAs that contain pause sites sustained, induced or resolved following the depletion of CNOT1. **B-D.** Examples of individual mRNAs with different Ccr4-Not regulated ribosome pause types. **B.** Pause resolved in the absence of CNOT1. **C.** Paused sustained in the presence and absence of CNOT1. **D.** paused induced by CNOT1 knockdown. Plots show RPF coverage normalised for mRNA abundance (TPM). **E.** The change in mRNA half-life after CNOT1 knockdown for the groups of mRNAs with different pause types identified in A. **F.** mRNAs with resolved pause sites in the absence of CNOT1 shown an increase in protein production relative to the ribosome occupancy on the mRNA when CNOT1 is depleted. **GHI.** Amino acid sequence motifs at the E, P, A sites of paused ribosomes and three codons upstream and downstream generated using Seq2Logo (113), shown for resolved pauses (G), sustained pauses (H) and induced paused (I).

It has been proposed that the Ccr4-Not complex can sense paused ribosomes and trigger mRNA decay (28), and examination of the change in mRNA half-life with CNOT1 knockdown of the distinct pause type groups shows mRNAs with sustained and induced pauses undergo greater stabilisation following CNOT1 depletion (Fig. 6E). This would fit with a model whereby the Ccr4-Not complex is involved in the sensing of stalled ribosomes and their resolution via decay mechanisms in that in the absence of CNOT1 the pauses are sustained or become pronounced.

In contrast, the mRNAs with pauses resolved by CNOT1 knockdown do not show an increase in mRNA half-life. To assess whether these are genuine sites of stalled ribosomes the unique combination of pulsed SILAC and ribosome profiling data was used. We observe that the ratio between protein production and ribosome occupancy increases on mRNAs with resolved pause sites after CNOT1 depletion (Fig. 6F). This demonstrates that CNOT1 knockdown resolves ribosome pauses on these messages leading to altered translation and protein synthesis.

Next, we examined whether there are specific sequence motifs associated with these Ccr4-Not complex regulated pause sites. Figures 6GHI show the amino acid sequence motif at the E, P and A-site and the 3 codons up and downstream for each of the pause site types (created using Seq2Logo (113)). This shows a strong enrichment for charged amino acids, particularly glutamate, being encoded at the A-site position of resolved and sustained pauses (Fig. 6GH). This suggests this is a specific motif for ribosome pausing in control conditions, but there may be additional factors that determine precisely how the Ccr4-Not complex acts upon the pause site. A sequence motif at induced pause sites is not so pronounced but has an enrichment for aspartate at the ribosome E-site (Fig. 6I).

Previously, ribosome pausing on two specific proteasome component mRNAs in yeast has been shown to be regulated by Not1 to facilitate co-translational assembly (12). Although there is still more to be elucidated about the precise mechanisms at play, this study now provides evidence for the role of the Ccr4-Not complex in the regulation of ribosome pausing in human cells and this regulation appears to be more widespread.

## Discussion

Numerous mechanisms for mRNA specific recruitment of Ccr4-Not complex exist, whereby sequence motifs, predominantly in the 3’UTR, are recognised by sequence specific RNA-binding proteins or microRNAs, resulting in the delivery of the Ccr4-Not complex to the mRNA (61,83,114–117). Importantly the Ccr4-Not complex has been shown to be able to exert both translational inhibition and mRNA destabilisation and that these effects can occur separately (39, 41).

The CNOT1 subunit of the Ccr4-Not complex functions as a scaffold protein bringing into proximity the core subunits, and regulatory proteins to coordinate the varied roles of the complex (5,118,119). CNOT1 also interacts with proteins such as TNRC6 that recruit the Ccr4-Not complex to the mRNA during miRNA-mediated repression (81, 114). Here we have depleted CNOT1 to examine the global effects upon mRNA stability (Fig. 1, Additional File 1: Fig. S1,2), the translational status of the mRNAs (Fig. 2, Additional File 1: Fig. S4,5), and protein production (Fig. 4B, Additional File 1: Fig. S8). This is the first such comprehensive investigation of this multifunctional protein complex and provides a benchmark dataset for translational studies. We demonstrate for the first time in human cells how the Ccr4-Not complex differentially regulates cohorts of mRNAs and how translational repression is distinct from how the complex regulates mRNA deadenylation and stability.

Codon usage, and more recently amino acid usage, have been associated with differences in mRNA stability in control conditions in multiple organisms (51,52,78,80,120–122). An example is reporter studies in yeast that have shown that substitution of optimal codons with synonymous non-optimal (generally A/U-ending) ones reduces mRNA stability (52). Recent studies in human cells have also found A/U-ending codons to be destabilising (80, 122), whereas another study has found G/C-ending codons to be destabilising (93). It is of note that this differential regulation may change with cellular context as it has been demonstrated that mRNAs preferentially involved in proliferation (enriched for A/U-ending codons) and differentiation (enriched for G/C-ending codons) have distinct codon usage and the tRNA pool available is altered to reflect this (76,77,123). Thus, the inconsistency in whether it is the A/U-ending or G/C-ending synonymous codons that are classified as destabilising in the literature could be explained by conditional differences.

Moreover, there is minimal understanding of the proteins involved in coordinating the response to codon usage. Previous research in yeast and zebrafish has implicated subunits of the Ccr4-Not complex in codon-mediated regulation of mRNA stability (16,28,72). We show that knockdown of CNOT1 preferentially stabilises mRNAs enriched in G/C-ending codons (Fig. 1DF, Fig. 5A) and thus show the central role of the Ccr4-Not complex in the link between mRNA stability and codon usage in human cells. Studies of the translatome and mRNA half-life are not often complemented with protein level data. Unexpectedly, we show that while large changes in mRNA stability attributable to codon composition of the CDS are observed (Fig. 1F,5A), these correlations are not apparent in the pulsed SILAC data (Additional File 1: Fig S9D).

Gene ontology analysis highlighted that increased mRNA half-life after CNOT1 knockdown is associated with an enrichment for biological processes relating to cardiac septum and muscle organ development (Additional File 1: Fig. S3A), which is particularly interesting given that recent publications demonstrate the importance of CNOT1 for cardiac development (68) and neurodevelopment (124). Decreased translational efficiency in the absence of CNOT1 is observed for mRNAs encoding proteins involved in extracellular structure organisation that preferentially localise to the ER and plasma membrane (Fig. 3, Additional File 1: Fig. S7). We also find that the reduced ribosome occupancy among this group of mRNAs following CNOT1 depletion occurs downstream of the signal sequence cleavage site (Fig. 3JK) and reduced localisation of mRNAs to the ER after CNOT1 depletion (Fig. 3H). An intriguing hypothesis would be that one of the translational control mechanisms mediated by the Ccr4-Not complex is coordination of subcellular localisation of mRNA.

In addition, our data shows a role for the Ccr4-Not complex in the translational repression of mRNAs that are localised to p-bodies (Additional File 1: Fig. S9C). It will be interesting to investigate in the future whether this is due to an indirect role of CNOT1 in p-body formation (96) or if the Ccr4-Not complex is also involved in mediating repression that occurs within these granules. The mRNAs translationally upregulated following CNOT1 depletion encode transcription factors/nuclear proteins (Fig. 5E, Additional File 1: Fig S10D), in agreement with p-body studies (93, 94). They are also enriched for amino acids associated with disordered regions in the proteins (Fig. 5C), intrinsically disordered regions have been suggested to be the regions of transcription factors that interact with the promoter region (125, 126). Also, the tertiary structure of zinc-finger domains has been shown to be able to act as a nuclear localisation sequence (127, 128), so perhaps the distinct amino acid composition of this group of mRNAs pertains to the localisation and protein function.

Finally, we identify groups of mRNAs on which ribosome pausing occurs in a CNOT1-dependent manner (Fig. 6A). These pauses impact how the Ccr4-Not complex regulates mRNA half-life (Fig. 6E) and protein synthesis (Fig. 6F). There is an enrichment for charged amino acids at the A-site of paused ribosomes in the presence of CNOT1 (Fig. 6GH). Whether there are additional proteins and/or sequence motifs that determine precisely how the Ccr4-Not complex regulates the fate of mRNAs with paused ribosomes could be investigated in the future.

## Conclusions

Here we have demonstrated that the CDS composition of an mRNA is important for the regulation of its fate by the Ccr4-Not complex in terms of codon usage and CDS length. In this cellular context G/C-ending codons mediate the destabilisation of an mRNA by the Ccr4-Not complex. We also discover a novel role for the Ccr4-Not complex in the regulation of the localisation of mRNAs to the ER for their translation. Moreover, mRNAs encoding proteins that localise to the nucleus are regulated at the level of translation by the Ccr4-Not complex and are sequestered in p-bodies in control conditions. Overall, we show that the Ccr4-Not complex is a control hub that governs multiple mechanisms to precisely regulate the fate of each mRNA.

## Supporting information

Supplemental Statistics

## Availability of data and materials

The datasets generated and/or analysed during the current study are available in the GEO and ProteomeXchange repository. Transcriptional inhibition experiments RNA-seq data have been deposited under GSE158619. Ribosome profiling experiments (both small RNA and total RNA libraries) have been deposited under GSE134517 for control siRNA and GSE158141 for siCNOT1 treated samples. SILAC data is deposited under PXD015772 (repeat 1) & PXD020305 (repeats 2 & 3). Code used for the analysis will be made available on github.

## Authors’ contributions

SG, AW and MB designed the experiments and wrote the manuscript. SG performed the ribosome profiling, transcriptional inhibition experiments, qPCR validations and all bioinformatic analysis. CG conducted ER/cyto fractionation experiments. AW, KH and SZ designed and conducted the pulsed SILAC experiments. All authors have approved the final manuscript.

## Experimental Methods

### Cell culture

HEK293 cells were cultured in Dulbecco’s Modified Eagle’s Medium (DMEM) supplemented with 1% L-glut and 10% FBS.

### siRNA treatment

Control siRNA (#3 Dharmacon) or CNOT1 siRNA (Ambion no.S22844) was transfected using DharmaFECT 1 (2:1 ratio of siRNA to DharmaFECT 1) to a final concentration of 30nM and cells harvested after 48hrs. Due to CNOT1 siRNA treatment causing slowed cell growth, cells for CNOT1 siRNA treatment were plated at 10% increased density to obtain the same cell numbers as the control samples at the time of harvesting. For experiments in Additional File 1: Fig. S2DEF & Fig. S9B an additional siRNA pool targeting CNOT1 was used (Horizon Discovery 015369-01)

### Antibodies

For western blot the antibodies used were: CNOT1 (ATLAS HPA Rabbit 046577, 1:500), vinculin (abcam mouse ab18058, 1in 10,000), GAPDH (CST 5174, 1:1000), Calnexin (CST 2679, 1:1000), rabbit secondary antibody (LI-COR Biosciences 926-32213, 1:10,000), mouse secondary antibody (LI-COR Biosciences 926-68072, 1:10,000).

### Ribosome Profiling

Ribosome profiling was conducted as previously described in Wilczynska et al 2019. In parallel to these control experiments, ribosome profiling was conducted for samples transfected with CNOT1-targeting siRNA for 48hrs. Three biological replicates were conducted.

### RT-qPCR from gradient fractions

For validation experiments, media was changed on a 15cm plate of HEK293 cells 1.5 hrs prior to harvesting. Cells were then scraped in ice-cold PBS, spun down and resuspended in lysis buffer containing cycloheximide. 300μl lysate was loaded on to a 10%-50% sucrose gradient then spun at 38000rpm for 2hrs at 4ᵒC. Gradient fractions were collected into 3ml 7M GuHCl, 8μl glycogen and 4ml ethanol added and then precipitated for >24hrs at -20ᵒC. The collected gradient fractions were pelleted at 4000 rpm for 1hr at 4ᵒC. The supernatant was removed, and the pellets resuspended in 400μl RNase-free water. The samples were then transferred to 1.5ml eppendorfs, 2μl glycogen, 40μl 3M NaOAc pH 5.2 and 1ml 100% ethanol added. These were precipitated overnight at -20ᵒC. Samples were then pelleted at 13000rpm 4ᵒC for 40 minutes, washed with 500μl 75% ethanol, air dried and resuspended in 30μl RNase-free water. To check the RNA integrity equal volumes (3ul) of each fraction along the gradient were ran on a 1% agarose gel.

RT-PCR was conducted on equal volumes (3ul) of each fraction using SuperScript III (Invitrogen 18080085). qPCR was then conducted using SYBR Green master mix (Applied Biosystems 4385618) on an Applied BioSystems QuantStudio 5 machine. For qPCR along the gradients fractions, the proportion of the mRNA present in fraction was plotted. All qPCR primers used are included in supplemental table 7.

### Cell viability

Cells were grown in 12-well plates and treated with a range of Triptolide concentrations. 100μμl trypsin was added per well and 200μl media added to quench this. 10μl of cells were then mixed with 10μl tryphan blue and cells negative and positive for tryphan blue counted using a haemocytometer.

### Transcriptional inhibition experiments

For the transcriptional inhibition experiments to determine mRNA half-lives, cells were plated in 12-well plates and transfected with control or CNOT1-targeting siRNA for 48hrs. The medium was changed 1 hr prior to treatment with 1μM triptolide (abcam: ab120720). At a range of time points (0, 0.5, 1, 2, 4, 8 & 16 hrs) post-triptolide addition, cells were washed with PBS and lysed directly in 1ml of Trizol for RNA samples or 150μl 1.5x SDS sample buffer for protein samples. Three biological replicates were conducted. The RNA was extracted with Trizol and acid-phenol chloroform. 3ug of RNA was then poly(A) selected (Lexogen 039.100). 4ng of poly(A) selected RNA was used as input into the CORALL Total RNA-Seq library prep kit (Lexogen 096.96) with 11 PCR cycles used. For qPCR validations (Additional File 1: Fig S2DEF) 1μM flavopiridol was used and 100ng/μl oligodT used in the RT reaction.

### Cytoplasmic/ER fractionation

Fractionation of cytoplasmic and ER material was performed by sequential detergent extraction, as previously reported (Reid, JBC, 2012). An isotonic buffer (20mM Tris pH 7.4, 150mM NaCl, 5mM MgCl_2_) was supplemented with 1mM DTT, 1x cOmplete EDTA-free protease inhibitor (Roche). For sequential lysis, it was further supplemented with 0.015% digitonin (cytosolic buffer), or 0.004% digitonin (wash buffer), or 2% n-Dodecyl β-D-maltoside (ER buffer). 80 U/ml of Ribolock RNase Inhibitor (Thermo Scientific) were added to each final buffer. 2/3 of each sample was process for RNA extraction with Trizol LS, and 5x SDS sample buffer added to the remaining lysate for western blotting.

Prior to library preparation ERCC spike-ins (Invitrogen) were added proportionally between the cytosolic and ER fraction in each condition. 900 ng of RNA was rRNA depleted with the RiboCop rRNA Depletion Kit HMR V2 (Lexogen) and library preparation performed with the CORALL Total RNA-Seq Library Prep Kit (Lexogen 096.96). Sequencing was performed on a NextSeq 550 system (Illumina).

### Pulsed SILAC (stable isotope labelling by amino acids in cell culture)

These experiments were conducted as described in Wilczynska et al 2019 (repeat 1 – forward and reverse - is the data used in Wilczynska et al 2019). In brief, HEK293 cells were cultured and siRNA treated for 30 hours. This media was then replaced with DMEM that does not contain Arginine or Lysine (Life Technologies). For the medium-heavy isotope containing medium, a supplement of [13C6] L-arginine (Arg-6) and [2H4] L-lysine (Lys-4) was added (Cambridge Isotope Laboratories). For the heavy isotope containing medium, [13C6][15N4] L-arginine (Arg-10) and [13C6][15N2] L-lysine (Lys-8) were added (Cambridge Isotope Laboratories). Both forward (heavy CNOT1 siRNA/medium-heavy control siRNA) and reverse (medium-heavy CNOT1 siRNA/heavy CNOT1 siRNA) replicates were conducted for each biological repeat. After 14 hrs cells were lysed in SDS-free RIPA buffer, pooled in a 1:1 ratio, reduced with DTT and alkylated with iodoacetamide. The samples were then trypsin digested and fractionated using reverse phase chromatography. For the exact details of the analysis of the samples by mass spectrometry and the data analysis with MaxQuant software (129) see Wilczynska et al 2019. A protein was retained for downstream analysis if it was detected in the forward and reverse replicate for a given biological repeat, and if detected in at least two of the three biological repeats.

### Luciferase reporter experiments

24hrs after siRNA transfection in 12-well plates, cells were transfected with 40ng pRL and 160ng pGL3 intron as a transfection control (Meijer NAR 2019) using 0.6ul GeneJammer (Agilent). The Renilla construct (pRL) was either the original sequence or a sequence with all the AUU/GUU/ACU codons converted to their synonymous codons (AUC/GUC/ACC). After another 24hrs, samples for detection of luciferase activity were washed twice with PBS and lysed in 1x passive lysis buffer, and 10µl lysate used for luciferase detection using the Dual Luciferase Reporter Assay System (Promega). Samples for RNA were harvested in 1ml Trizol. Relative luciferase activity was determined by the ratio between Renilla and Firefly luciferase and the relative luciferase RNA-levels determined by qPCR. The translational efficiency of Renilla was then determined by luciferase activity / RNA-level.

## Data Analysis Methods

### mRNA half-life experiments RNA sequencing data processing

Cutadapt (130) was used to removed adapters. cd-hit-dup (131) was used to deduplicate the reads based on the 12nt long UMIs. The remaining reads were aligned to the genome using STAR (132) and a gtf file filtered to contain the most abundant transcript per gene. To obtain the read counts featureCounts (133) was used with the gtf file filtered for the most abundant transcript per gene. The read counts were first normalised for the library size. As the libraries are prepared based on equal ng of material, the read counts are normalised back to the nanodrop concentrations of the RNA that was obtained from equal cell numbers. For each condition the data was then normalised relative to the 0hr time point to allow for comparison between mRNAs and conditions.

### Modelling mRNA decay rate

To be able to use the three replicates together in the decay modelling, for each replicate the values for each mRNA across the time points were normalised to the 0hr time point (set at 100). The simple model for mRNA decay is that it follows the exponential decay function: y ∼ y_0_e^-kt^. Where y_0_ is the steady state mRNA level, k is the decay constant and t is time. Outliers were first identified based on the methodology described in (134). The nlrob function (R package: robustbase) was used to fit a robust nonlinear model to the data with an adapted form of the NLS.expoDecay function (R package: aomisc) to provide start parameters for the model fit. To next identify possible outliers the weighted was calculated: weighted residual = absolute(observed – expected)/expected. The maximum proportion of outliers was set to 20% to ensure natural biological variation was not mistaken for an outlier. The robust standard deviation of residuals (RSDR; (134)) was then calculated by ranking the residuals in terms of absolute value and taking the value at the 68.27 percentile and multiplying this value by N/(N-K), where N is the number of values and K is the number of parameters being modelled. The significance level for outlier removal is α_i_ = Q(N-(i-1))/N, where N is the number of values in the data and i is the ith value in the ordered list of residuals. Q was set to 5% (0.05) and this means that there is a 5% chance of falsely discovering a significant outlier. The t-score in this case in then calculated by: t-score = residual_i_/RSDR (134). The pt function (R package: stats) is then used to obtain the two-tailed p-value of the t-score. If this p-value is less than α_i_ then this value is a significant outlier.

Once significant outliers had been removed, the modelling was conducted in R using the nls function from the stats package. To ensure appropriate starting parameters were used in the modelling a self-starting function was used – NLS.expoDecay() part of the aomisc R package. The half-lives were then calculated from the decay rate using the equation: t_1/2_=ln(2)/k.

### Assessment of feature influence

To assess which mRNA features contribute to differential regulation of mRNA fate by the Ccr4-Not complex a supervised learning approach of gradient boosting (gbm R package) was used (Fig. 1D & 4C). Only one of highly correlated features (r > 0.7) were retained for the analysis (Additional File 1: Fig. S3B). For Fig. 1D a gaussian distribution was assumed and for Fig. 4C a multinomial distribution. The parameters used were: n.trees = 200, interaction.depth = 6, shrinkage = 0.005, cv.folds = 10.

### Ribosome profiling - Small RNA alignment and counts

For the small RNA sequencing data (ribosome protected fragments: RPFs), Cutadapt (130) was used to remove the adapter sequence and the reads were deduplicated based on the 8nt unique molecular indexes (UMIs - 4nt either end of the read) using cd-hit-dup (131). Cutadapt (113) was then used to remove UMIs and to select read lengths 25nt to 35nt (the expected size range of the RPFs). The reads were first aligned to a fasta file of rRNA sequences, to remove contaminant rRNA fragments by alignment with bowtie (135). The reads were then mapped with bowtie to the hg38 gencode version 28 (136)) protein coding transcriptome that had been filtered for the most abundant transcript per gene as determined from the control total RNA-seq data.

To get the number of RPFs per gene and the exact position of the RPFs along the mRNA a python script from the RiboCount part of the RiboPlot package (https://pythonhosted.org/riboplot/) was adapted. This was first conducted for each read length to determine the frame, periodicity and P-site offset for each read length. It was determined that read lengths 27 to 31 showed strong RPF characteristics, thus these read lengths were selected for downstream analysis. E,P & A-site offset were determined to be 9nt, 12nt & 15nt from the read start respectively (see Additional File 1: Fig. S11AB). For figures of ribosome position across individual mRNAs the P-site position was used (Fig. 6BCD, Additional File 1: Fig.S6).

### Ribosome Profiling - Total RNA alignment and counts

For the corresponding total mRNA samples, cutadapt (130) was used to remove adapter sequences, cd-hit-dup (131) to deduplicate based on the 8nt UMI. The UMI was then removed with cutadapt and the sequences aligned with STAR (132) to a gtf file filtered for the most abundant transcript per gene. Bam files were sorted and indexed using SAMtools (137). Read counts were obtained using htseq-count (138).

### DESeq2 differential expression analysis

DESeq2 (86, 87) was used for differential expression analysis of the RNA and RPFs following CNOT1 knockdown. As the RPFs and RNA are very different library types, something the DESeq2 package is not able to account for, the differential expression analysis was conducted independently for the two datasets. The data was pre-filtered to ensure at least three samples had a minimum of 10 read counts for any given mRNA. To ensure that lower abundance mRNAs or lowly translated mRNAs do not have an exaggerated fold change the lfcShrink function with apeglm model of the DESeq2 package was used for effect size estimations (87). Log2FC translational efficiency was calculated as log2FC RPF – log2FC RNA from the DESeq2 results (Supplemental Table 2). DESeq2 was also used in the same manner to determine the log2FC in RNA in cytosolic and ER fraction following CNOT1 depletion (Supplemental Table 4).

### Differential ribosome occupancy across the CDS

For the total RNA data transcripts per million (TPM) was calculated. For the RPFs read the data was normalised for library size and for each read the nucleotide of the P-site start used (12nt offset from read start). RPF counts at each position along the CDS of each mRNA were then normalised for the mRNA abundance using the TPM from the total RNA-seq. After this normalisation to account for mRNA abundance changes, for each biological replicate a delta was conducted for the normalised RPF coverage along each mRNA between CNOT1 and control siRNA conditions. The delta for each mRNA was then averaged for the three biological replicates. For Fig.3EJ the delta in the distinct groups of mRNAs was binned into 40 windows across the CDS to account for the different CDS lengths of the mRNAs and the median change in ribosome occupancy in each window displayed.

### Gene Set Enrichment Analysis

Normalised enrichment scores for biological process, cellular component and molecular function gene sets were calculated using the fgsea R package. Significant enrichment was determined using an adjusted p-value threshold of <0.05. The full list of significant results is in Supplemental Table 3.

### Synonymous codon usage

Synonymous codon usage is based on the fact that for many amino acids there are multiple codons that encode it. For each codon the number present within a given CDS were counted and then normalised for the total number of possible codons that can encode the same amino acid in that CDS. This was conducted for each mRNAs and then for the distinctly regulated groups of mRNAs this was then averaged across the group. The heatmaps are column-scaled as the comparison being made is the preferential use of the codons between the different groups of mRNAs (Fig. 5A). The coloured bar indicates the nucleotide at the third position of the codon.

### Amino acid usage

For amino acid usage the frequency of codons for each amino acid were counted per mRNA and normalised for the number of codons in the mRNA. The frequency for each amino acid was then averaged across the group of mRNAs (if the amino acid frequency for an mRNA was zero it was excluded from the average). The heatmaps are column-scaled as the comparison being made is the use of the amino acid between the different groups of mRNAs (Fig. 5B). The coloured bar indicates the type of amino acid side chains. For Fig. 5CD amino acids were classified as disorder-promoting (P|Q|E|S|K) as in (108).

### Ribosome pause site determination

For analysis of paused elongating ribosomes all RPFs apart from those located in the first 15 and last 5 codons of the CDS were used. A pause site in each condition was defined as a position with a RPF peak height ten times greater than the average RPF peak on the mRNA. The change in peak height between conditions was determined as a delta between RPFs that had been normalised for the mRNA abundance (TPM). Pause sites were then classified as either ‘sustained’ if there was no change in peak height; ‘resolved’ if RPF peak height decrease was ten times greater than the average delta decrease across the mRNA or ‘induced’ if the increase in RPF peak height was ten times greater than the average delta increase (Fig. 6A). The mRNAs distinctly with one type of pause site are used in downstream analysis and the exact pause positions are indicated in Supplemental Table 6.

### Motif analysis

For the amino acid motifs generated in Fig .6GHI the Seq2Logo web app was used (113). The settings used were P-Weighted Kullback-Leibler logo type, Hobohm1 clustering method and 200 weight on prior.

### k-means clustering

For clustering of mRNAs based on their half-lives (Fig. 1B) in the presence and absence of CNOT1 the half-life values were log transformed and the optimal number of clusters determined using within cluster sum of squares. The factoextra R package was used for kmeans analysis and cluster visualisation. For clustering of mRNAs into ER-targeted and cytosolic mRNAs in control conditions (Fig. 3C) the RNA-seq counts were first transformed into counts per million (CPM) and adjusted using the ERCC spike-in sequences to account for differences in absolute RNA levels between the cytosolic and ER fractions.

### Statistics

The Kruskal Wallis test was used for Figure 3FGHI. For all the figures requiring multiple comparisons the dunn test (FSA R package) was used to determine significance with Benjamini Hochberg used to correct for multiple hypothesis testing. For the groups of mRNAs distinctly regulated by the Ccr4-Not complex as identified in Fig. 4A the full set of statistical comparisons between the data for each group are included in tables in Additional File 2 for clarity. *** indicates p.adj<0.001, ** p.adj<0.01 & * p.adj<0.1. For Additional File 1: Fig S9B a two-tailed paired t-test was used.

## Additional File 1: Supplemental Figure Legends

**Fig. S1: Transcriptional inhibition experiment quality control. A.** Change in the synthesis of Ccr4-Not complex subunits following CNOT1 depletion, data from the pulsed SILAC experiments detailed in Additional File 1: Fig. S8. ND = not detected. **B.** Triptolide does not negatively impact cell viability in the time period used. Cell viability was assessed using tryphan blue staining at a range of 0-10μM triptolide after 16 hrs treatment. **C.** RNA integrity from triptolide-treated cells was assessed by agarose gel. **D.** qPCR for MYC across the range of triptolide concentrations at 4, 8, and 16hrs post-treatment shown relative to the MYC level at 0 hrs. **E.** Representative western blot confirming CNOT1 knockdown is maintained throughout the triptolide treatment. **F.** Representative agarose gel showing RNA integrity across the timepoints and conditions.

**Fig. S2: Validation of mRNA half-lives. ABC.** Example mRNA half-lives from the sequencing data used to determine global mRNA half-lives in the presence and absence of CNOT1. Examples are included for two mRNAs in each of the clusters in Fig. 1B. **DEF.** qPCR validations for the same mRNAs in ABC but using an alternative transcriptional inhibitor flavopiridol and an additional pool of siRNAs targeting CNOT1. This validates the mRNA half-lives for each of the clusters identified in Fig. 1B **D.** cluster 1. **E.** cluster 2 and **F.** cluster 3.

**Fig. S3: mRNA feature analysis. A.** The significant (adj.pval < 0.05) gene ontology biological process terms associated with an increase in mRNA half-life following CNOT1 depletion (conducted using the fgsea R package). **B.** Correlation matrix to identify highly correlated features. For features with a correlation coefficient > 0.75 only one of the features was retained. Retained features taken forward to feature importance analysis are highlighted. **C.** The correlation between the GC content of the CDS and the GC content of the 3^rd^ nucleotide of the codons in the CDS. **D.** Correlation coefficient (Spearman’s Rho) between the frequency of a given codon and the log2FC mRNA half-life (siCNOT1 / siControl) as in Fig. 1E, but reordered by the amino acid. Codons with an A/U at the 3^rd^ nucleotide position are coloured in magenta and codons with a G/C at the 3^rd^ nucleotide position are coloured in cyan. Amino acids are coloured by their charge/polarity type.

**Fig. S4: Ribosome profiling experiment quality control. A-C.** Representative examples of experimental quality control during RPF library production**. A.** TBE-Urea gel for extraction of RPFs (left=before, right= after extraction). Incision is made inclusive of the 28nt and exclusive of the 34nt marker position to avoid major contaminant rRNA fragments. **B.** Small RNA bioanalyzer chip traces of the extracted fragments before and after rRNA removal with Illumina RiboZero. **C.** TBE gel containing final small RNA libraries.

**Fig. S5: Ribosome profiling data quality control. A.** Proportion of reads that align to rRNA sequences in all samples. **B.** There is a high correlation across the biological replicates for the RPF sample read counts in both the control siRNA and CNOT1 siRNA treated samples. Pearson’s r values are indicated. **C.** RPF samples show the expected read length distribution across the three replicates and both conditions. **D.** The majority of RPFs align to the CDS. Graph shows the proportion of RPFs aligning to the 5’UTR, CDS and 3’UTR. **E.** The frame distribution of read lengths 27 to 32 in control and CNOT1 siRNA treated samples. Data shown is for a representative replicate.

**Fig. S6: Ribosome occupancy for individual mRNA examples. A-F.** Average RPF read count from three biological replicates along the CDS of example mRNAs. This is shown for control (grey) and CNOT1 knockdown (coloured). Validation of ribosome profiling data (Fig. 2C) using qPCR along gradient fractions from an independent experiment with and without CNOT1 depletion (n=2, n=1 shown in Fig.2D-F). **AB.** POLB and HDHD3 that show increased TE after CNOT1 knockdown shift into polysomes. **CD.** DENR and INTS7 with no TE change after CNOT1 knockdown show minimal change in distribution across the gradient. **EF.** GIT1 and POLR3E that have decreased TE after CNOT1 knockdown shift towards the sub-polysomes.

**Fig. S7: Gene set enrichment analysis. AB.** All mRNAs were ordered by the log2FC TE (siCNOT1/siControl) and gene set enrichment analysis conducted on the ranked list using the fgsea R package for **A.** biological processes & **B.** Molecular function. Red highlights terms with an enrichment associated with increased TE and blue highlights terms linked with a decreased TE following CNOT1 depletion. Up to a maximum of 10 significant terms (adj.pval < 0.05) are shown in this figure, the complete list of results are in Supplemental Table 3. **CD.** LOPIT data (from U2OS cells, (88)) was used to examine where the encoded proteins localise. mRNAs with decreased TE following CNOT1 depletion are enriched for proteins that localise to **C.** the ER & **D.** the plasma membrane.

**Fig. S8: Pulsed SILAC quality control. A.** For each biological replicate of pulsed SILAC, two technical replicates are required – forward and reverse labelling. Light, medium-heavy and heavy indicate the isotope of lysine and arginine present in the medium. Cells were treated with control siRNA or CNOT1-targeting siRNA for 30 hrs, before addition of medium-heavy/heavy media to cells for 14hrs. **B.** Correlations between forward and reverse labelling methods and biological replicates for pulsed SILAC data. Shown is the Pearson correlation coefficient. **C.** Venn diagram of the overlap of proteins detected in each replicate. For each replicate only proteins detected in both the forward and reverse labelling were used.

**Fig. S9 Codon frequency correlation with log2TE. A.** Correlation coefficient (Spearman’s Rho) between the frequency of a given codon in an mRNA and the change in translational efficiency (siCNOT1/siControl) as determined by the ribosome profiling experiments. Codons with an A/U at the 3^rd^ nucleotide position are coloured in magenta and codons with a G/C at the 3^rd^ nucleotide position are coloured in cyan. **B.** The codons identified as most negatively correlated with the log2FC TE in (A) – AUC/GUC/ACC were introduced at synonymous positions in the Renilla CDS (AUU/GUU/ACU). Renilla translational efficiency is calculated by the luciferase activity / luciferase RNA level (after normalisation to Firefly luciferase transfection control). The graph shows the translational efficiency change with CNOT1 depletion and the difference between the original Renilla CDS and the codon altered CDS. **C.** mRNAs with increased TE when CNOT1 is depleted are enriched in p-bodies in control conditions (data from HEK293 cells, (94)). **D.** Correlation coefficient (Spearman’s Rho) between the frequency of a given codon in an mRNA and the change in protein production (siCNOT1/siControl) as determined by pulsed SILAC experiments. Codons with an A/U at the 3^rd^ nucleotide position are coloured in magenta and codons with a G/C at the 3^rd^ nucleotide position are coloured in cyan.

**Fig. S10: mRNAs only upregulated translationally after CNOT1 depletion encode nuclear proteins. A.** Diagram to illustrate the categorisation of mRNAs in Fig. 4A by how the Ccr4-Not complex differentially impacts the translation and/or stability of the mRNA **B.** Distribution of 3’UTR lengths in the groups of mRNAs differentially regulated by the Ccr4-Not complex as classified in Fig. 4A. All comparisons were determined to be statistically significant using the dunn test and are shown in Additional File 2: Fig. S2B. **C.** The 3’UTR A content for the distinct groups of mRNAs. Statistical comparisons for A & B are shown in Additional File 2: Fig. S2BC. Only the non-significant comparisons are displayed for clarity and all statistics are presented in Additional File 2: Fig. S2B. **D.** Percentage of the groups of mRNAs in Fig.4A that localise to the nucleus/chromatin, determined using LOPIT data (from U2OS cells, (88)). **E.** Example mRNA stability profiles with control or CNOT1-targeting siRNA treatment for mRNAs in the TE ↑ group (dark red, Fig. 4A). They have a short half-life in control conditions that does not change with CNOT1 knockdown.

**Fig. S11: Precise ribosome residency. A.** To be able to assign the location of the E/P/A tRNA binding sites of the ribosome, an offset needs to be applied from the start of the read. The start codons have high ribosome occupancy so by looking at the positions of the read starts around this region, as in B, the offset to position the peak at the P-site. **B.** Plots show the RPF read distribution around the AUG start codon of all transcripts for a representative replicate. P-site offset is determined to be 12nt and thus the E-site offset is 9nt and A-site offset 15nt.

## Additional File 2: Statistics

**Fig. S1: Statistics for Main** **Figure 4**. Dunn test was used with Benjamini-Hochberg correction for multiple hypothesis testing correction.

**Fig. S2: Statistics for Main** **Figure 5C** **Additional File 1 S10BC.** Dunn test was used with Benjamini-Hochberg correction for multiple hypothesis testing correction.

## Supplementary Tables

**Table 1:** log2FC mRNA half-life

**Table 2:** log2FC RNA & log2FC RPF (DESeq2 data)

**Table 3:** Gene Ontology fgsea results

**Table 4:** ER/cyto fractionation RNA-seq

**Table 5:** pulsed SILAC

**Table 6:** pause site types

**Table 7:** qPCR primer sequences

