## Supplemental Figures for "Differential regulation of mRNA fate by the human Ccr4-Not complex is driven by CDS composition and mRNA localisation"

Fig. S1

A.

| Complex subunit | log2FC protein production |
|---|---|
| CNOT1 | -3.10 |
| CNOT2 | -0.23 |
| CNOT3 | 0.29 |
| CNOT4 | 0.09 |
| CNOT6/6L | ND |
| CNOT7 | -0.82 |
| CNOT8 | ND |
| CNOT9 | -1.64 |
| CNOT10 | -0.98 |
| CNOT11 | -0.11 |
| TAB182 | 0.46 |

B.

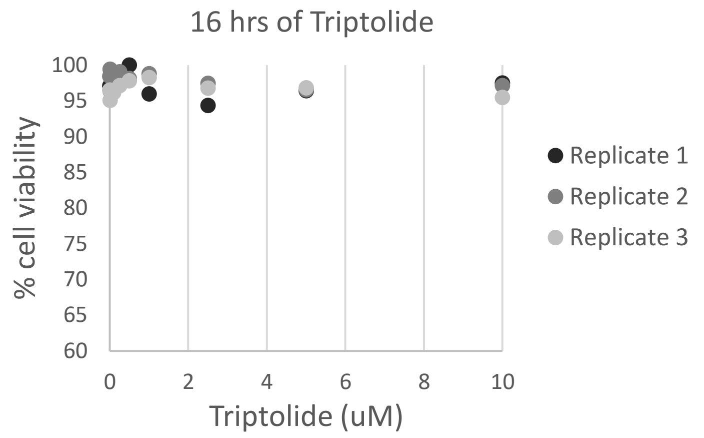


C.

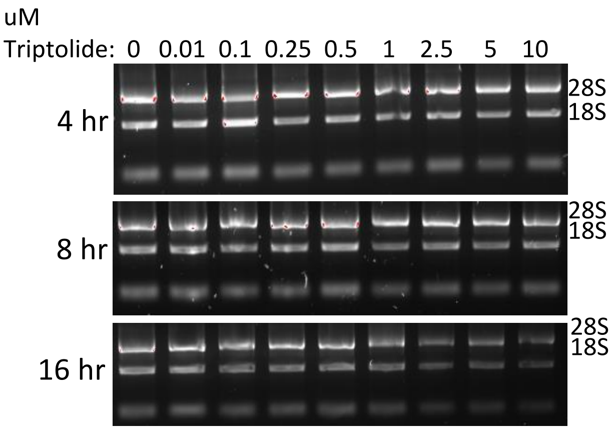


D.

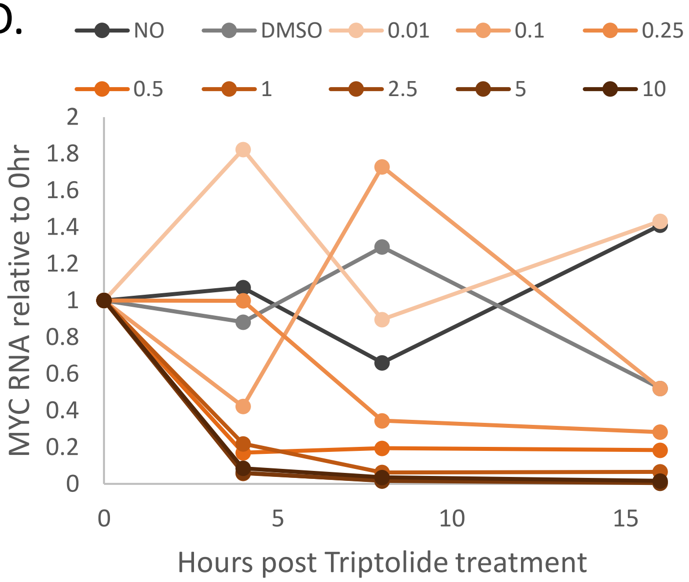


E.

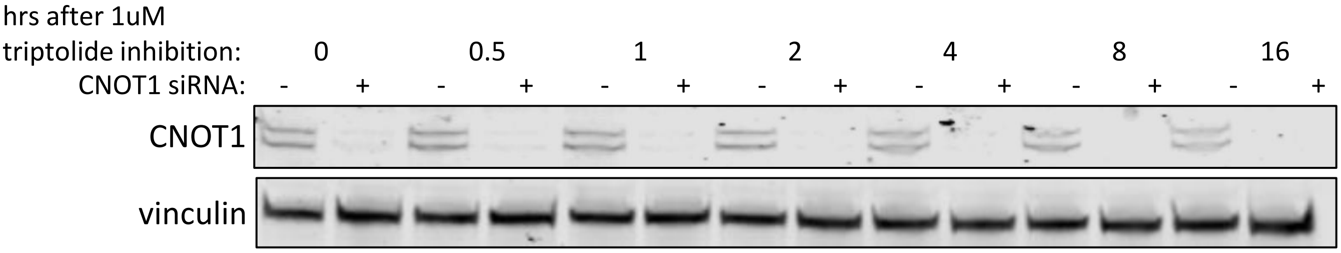


F.

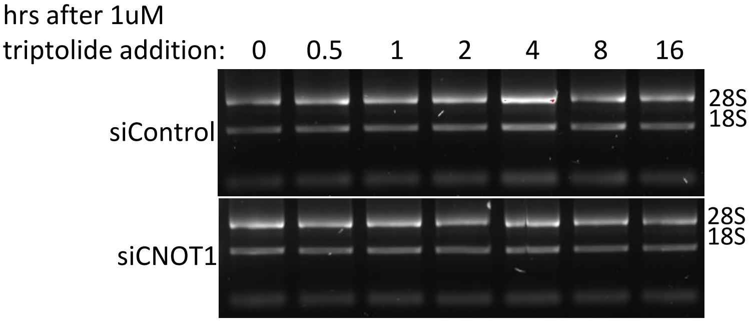

Fig. S2

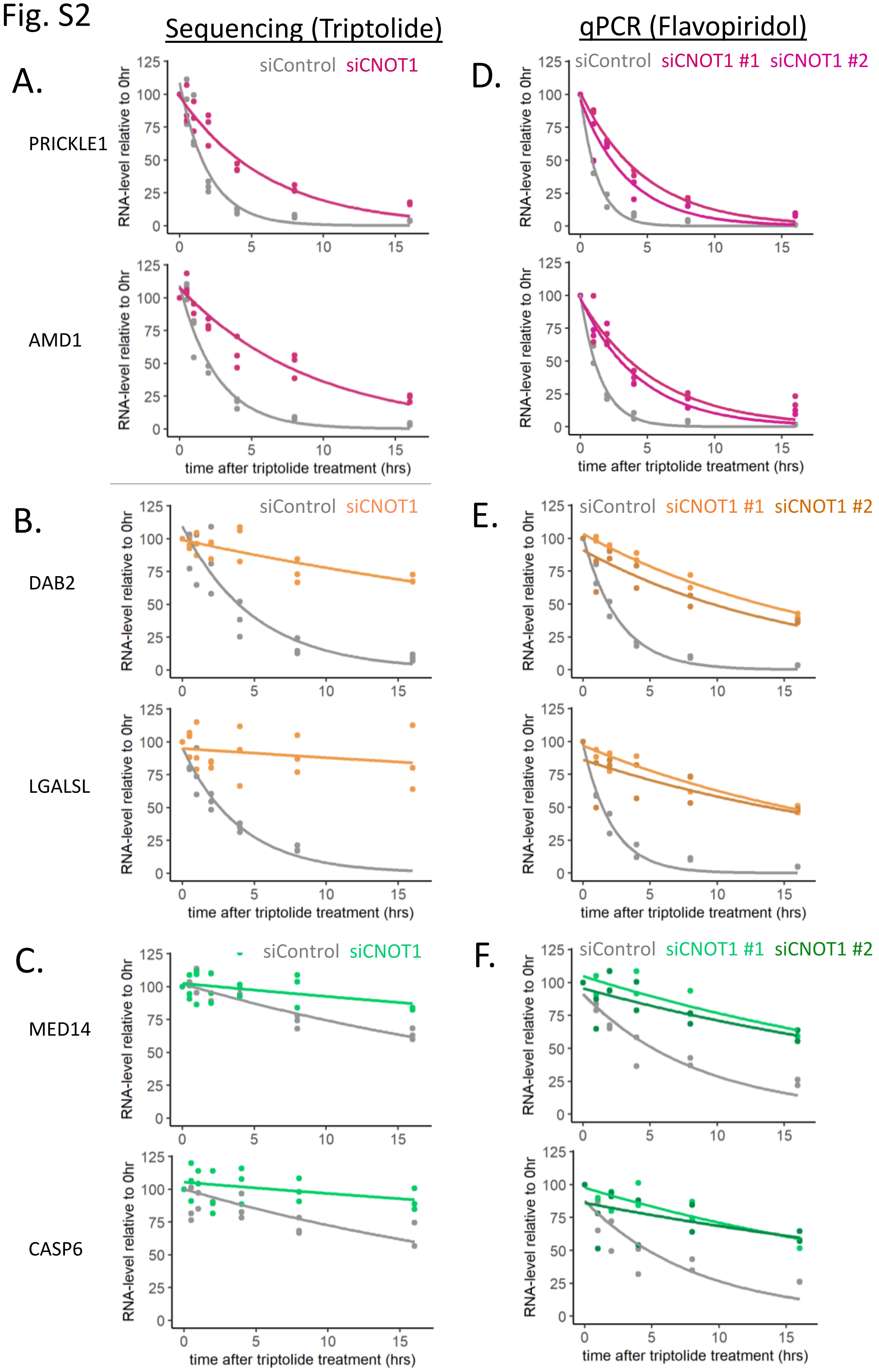

Sequencing (Triptolide)
qPCR (Flavopiridol)
A.
PRICKLE1
AMD1
siControl
siCNOT1
D.
siCNOT1 #1
siCNOT1 #2
B.
DAB2
LGALSL
E.
C.
MED14
CASP6
F.
RNA-level relative to 0hr
time after triptolide treatment (hrs)

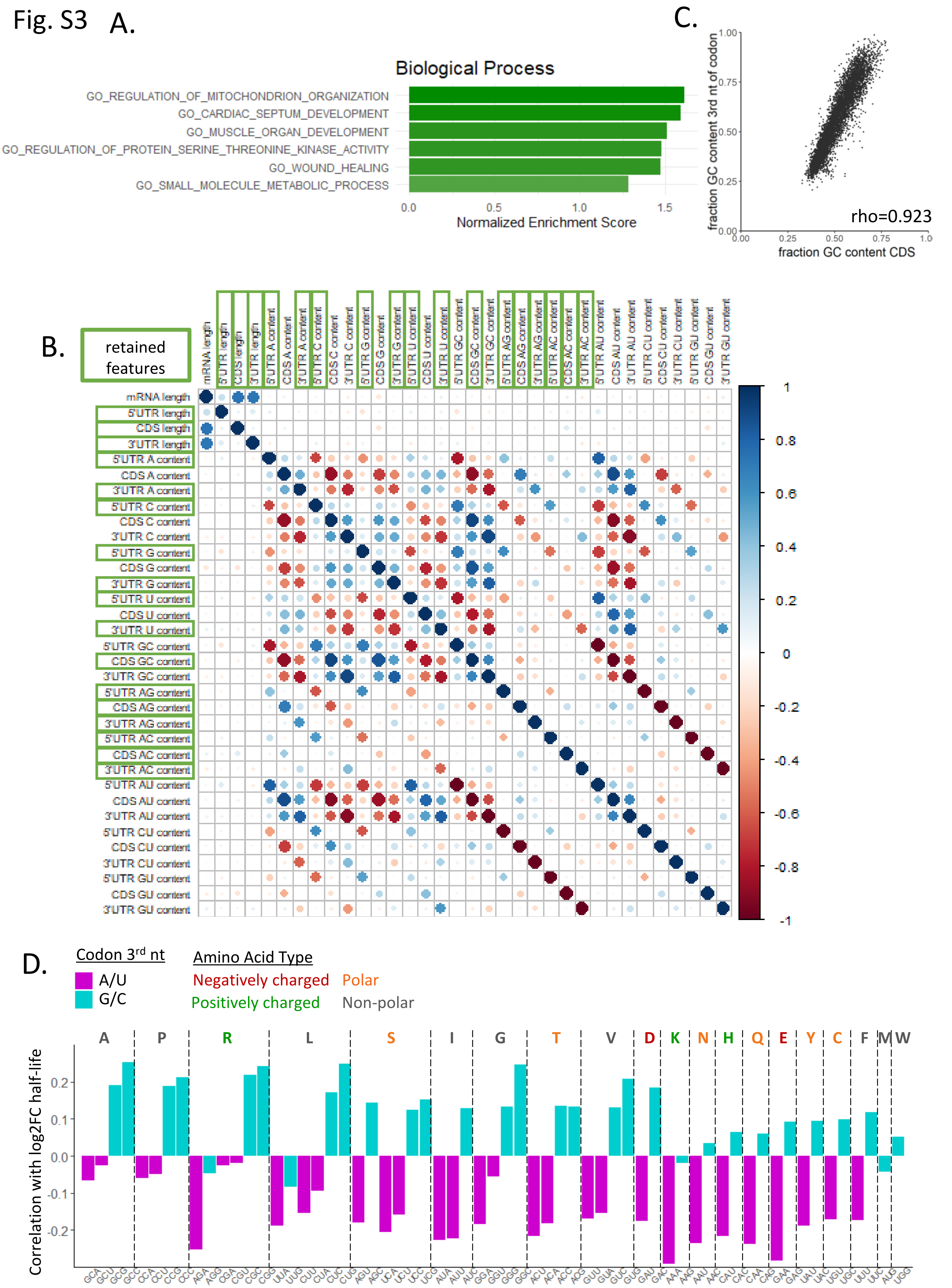
A.
Biological Process
GO_REGULATION_OF_MITOCHONDRION_ORGANIZATION
GO_CARDIAC_SEPTUM_DEVELOPMENT
GO_MUSCLE_ORGAN_DEVELOPMENT
GO_REGULATION_OF_PROTEIN_SERINE_THREONINE_KINASE_ACTIVITY
GO_WOUND_HEALING
GO_SMALL_MOLECULE_METABOLIC_PROCESS
0.0
0.5
1.0
1.5
Normalized Enrichment Score
C.
fraction GC content 3rd nt of codon
fraction GC content CDS
0.00
0.25
0.50
0.75
1.00
rho=0.923
B.
retained features
mRNA length
5'UTR length
CDS length
3'UTR length
5'UTR A content
CDS A content
3'UTR A content
5'UTR C content
CDS C content
3'UTR C content
5'UTR G content
CDS G content
3'UTR G content
5'UTR U content
CDS U content
3'UTR U content
5'UTR GC content
CDS GC content
3'UTR GC content
5'UTR AG content
CDS AG content
3'UTR AG content
5'UTR AC content
CDS AC content
3'UTR AC content
5'UTR AU content
CDS AU content
3'UTR AU content
5'UTR CU content
CDS CU content
3'UTR CU content
5'UTR GU content
CDS GU content
3'UTR GU content
1
0.8
0.6
0.4
0.2
0
-0.2
-0.4
-0.6
-0.8
-1
D.
Codon 3rd nt
A/U
G/C
Amino Acid Type
Negatively charged
Polar
Positively charged
Non-polar
A P R L S I G T V D K N H Q E Y C F M W
Correlation with log2FC half-life
0.2
0.1
0.0
-0.1
-0.2
GCA GCU GCG GCC CCA CCU CCG CCC AGA AGG CGA CGU CGC CGG UUA UUG CUU CUA CUC CUG AGU AGC UCA UCU UCC UCG AUA AUU AUC GGA GGU GGG GGC ACU ACA ACC ACG GUU GUA GUC GUG GAU GAC AAA AAG AAU AAC CAU CAC CAA CAG GAA GAG UAU UAC UGU UGC UUU UUC AUG UGG

Fig. S3

Fig. S4

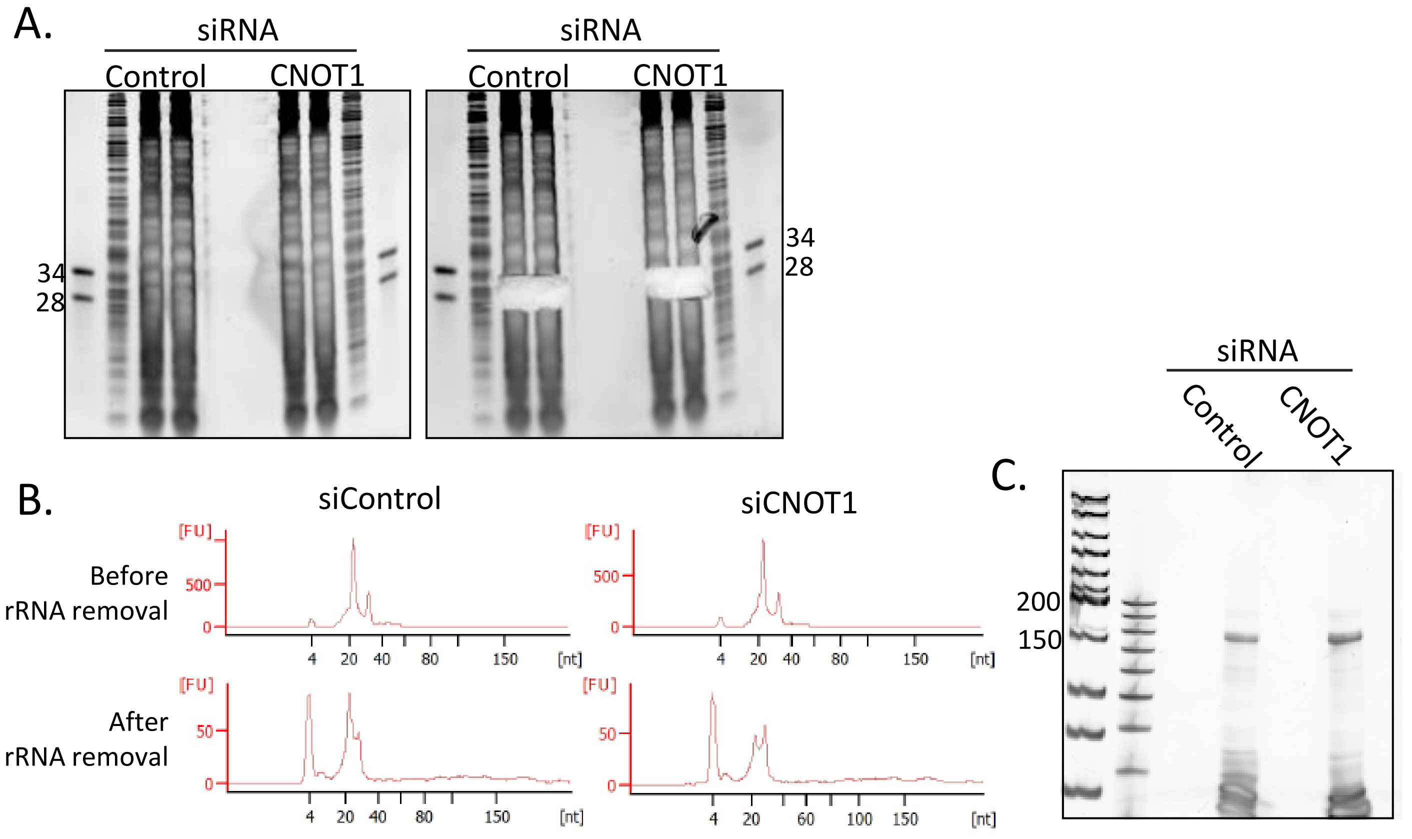
A.
siRNA
Control
CNOT1
34
28
siRNA
Control
CNOT1
34
28
B.
siControl
siCNOT1
Before
rRNA removal
After
rRNA removal
[FU]
500
0
50
4
20
40
60
80
100
150
[nt]
C.
siRNA
Control
CNOT1
200
150

Fig. S5

A.

| siRNA | Replicate | % rRNA |
|---|---|---|
| Control | 1 | 4.98 |
| Control | 2 | 5.00 |
| Control | 3 | 4.12 |
| CNOT1 | 1 | 11.05 |
| CNOT1 | 2 | 6.43 |
| CNOT1 | 3 | 8.40 |

B.

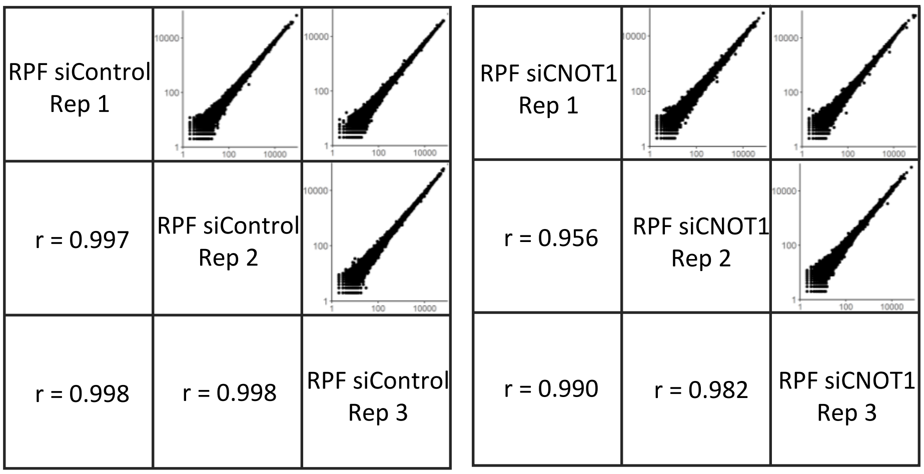


C.

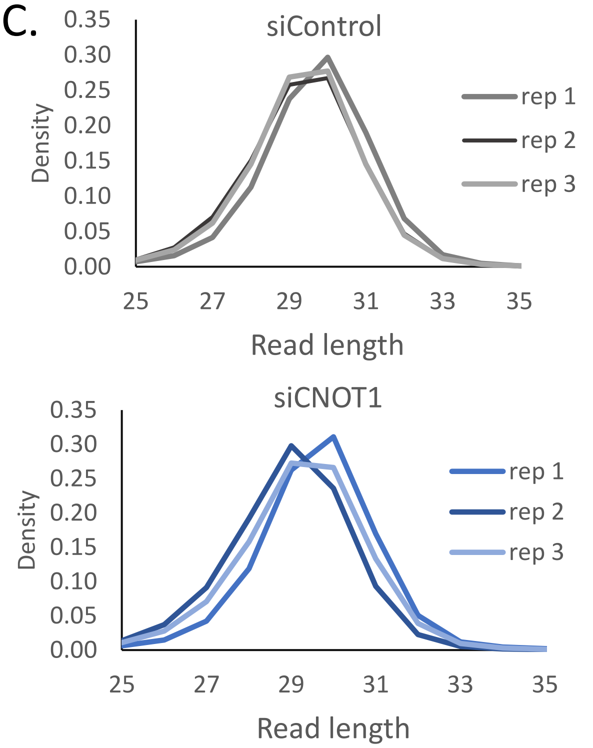


D.

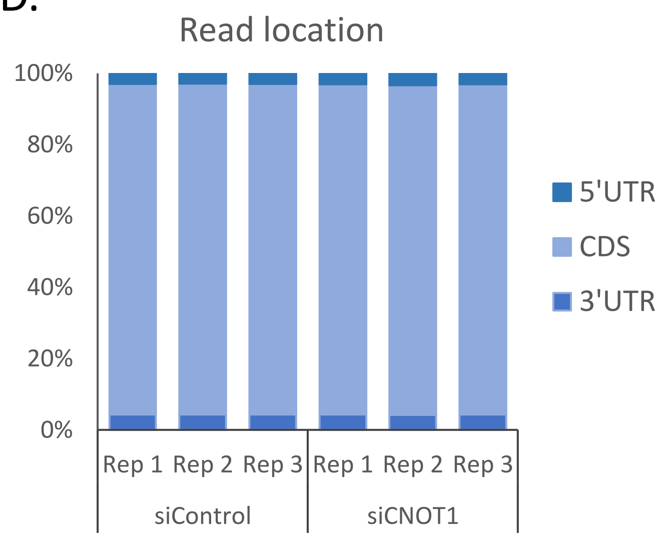


E.

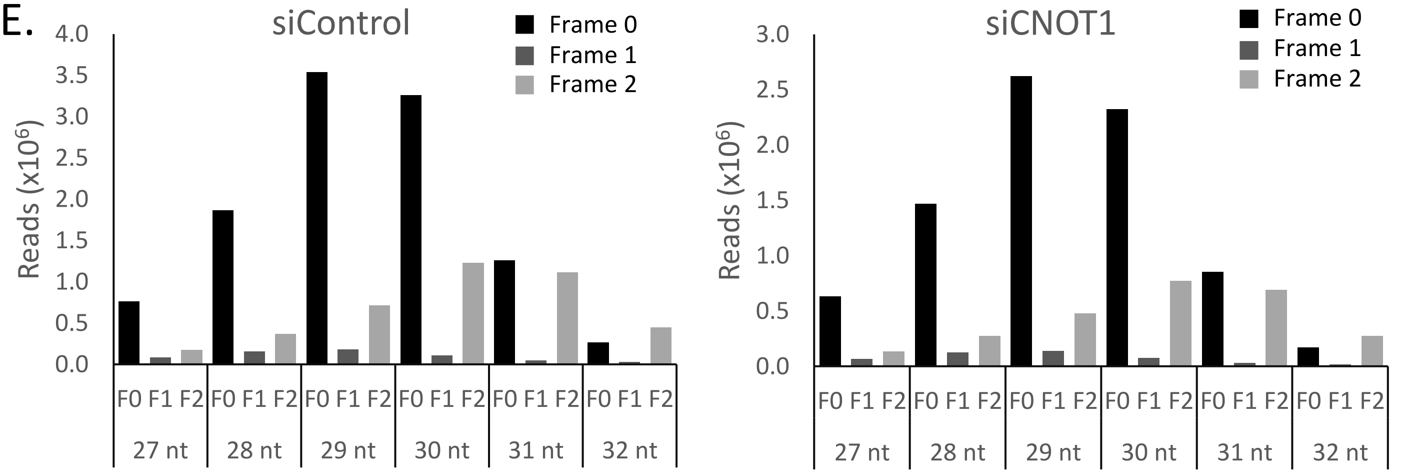

Fig. S6

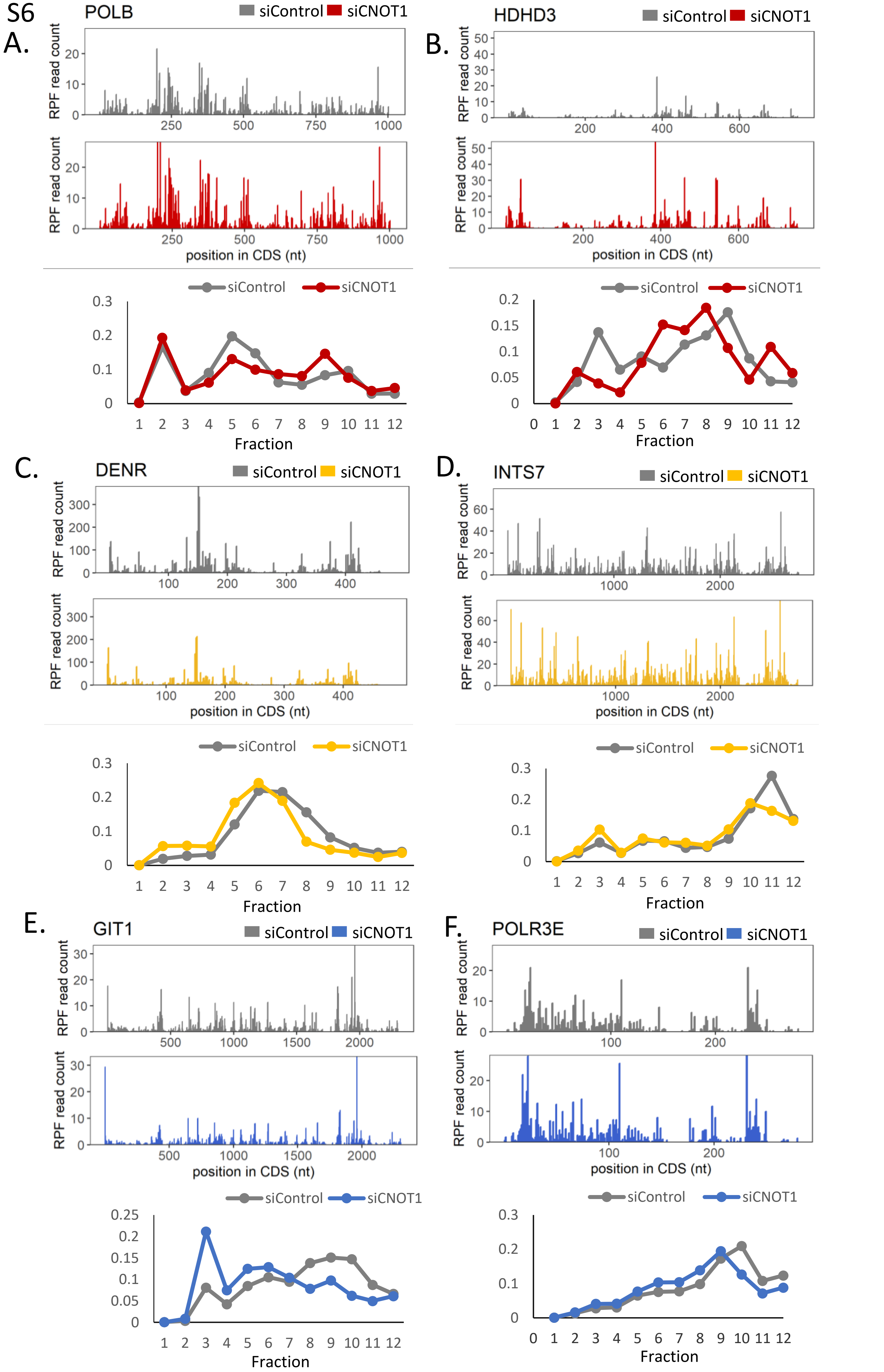
A.
POLB
siControl
siCNOT1
RPF read count
position in CDS (nt)
Fraction
B.
HDHD3
C.
DENR
D.
INTS7
E.
GIT1
F.
POLR3E

Fig. S7

A.

| Biological Process | Normalised Enrichment Score |
|---|---|
| NADH DEHYDROGENASE COMPLEX ASSEMBLY | 1.94 |
| MITOCHONDRIAL RESPIRATORY CHAIN COMPLEX ASSEMBLY | 1.90 |
| TRNA METABOLIC PROCESS | 1.75 |
| RESPIRATORY ELECTRON TRANSPORT CHAIN | 1.68 |
| REGULATION OF CENTROSOME DUPLICATION | 1.67 |
| TRNA MODIFICATION | 1.67 |
| REGULATION OF SISTER CHROMATID SEGREGATION | 1.66 |
| CENTROSOME DUPLICATION | 1.65 |
| MITOCHONDRIAL GENE EXPRESSION | 1.63 |
| TRNA PROCESSING | 1.62 |
| CARDIAC CONDUCTION | -2.00 |
| VASCULAR ENDOTHELIAL GROWTH FACTOR RECEPTOR SIGNALING PATHWAY | -2.01 |
| NEURON PROJECTION GUIDANCE | -2.03 |
| SECOND MESSENGER MEDIATED SIGNALING | -2.03 |
| REGULATION OF RESPONSE TO WOUNDING | -2.04 |
| REGULATION OF SYNAPSE STRUCTURE OR ACTIVITY | -2.05 |
| SYNAPSE ORGANIZATION | -2.05 |
| CYCLIC NUCLEOTIDE MEDIATED SIGNALING | -2.07 |
| CELL CELL ADHESION VIA PLASMA MEMBRANE ADHESION MOLECULES | -2.11 |
| EXTRACELLULAR STRUCTURE ORGANIZATION | -2.22 |

B.

| Molecular Function | Normalised Enrichment Score |
|---|---|
| TRNA BINDING | 1.77 |
| DNA BINDING TRANSCRIPTION REPRESSOR ACTIVITY | 1.63 |
| PASSIVE TRANSMEMBRANE TRANSPORTER ACTIVITY | -2.00 |
| ACTIVE ION TRANSMEMBRANE TRANSPORTER ACTIVITY | -2.00 |
| GATED CHANNEL ACTIVITY | -2.03 |
| PROTEIN KINASE ACTIVITY | -2.03 |
| STRUCTURAL MOLECULE ACTIVITY | -2.03 |
| MOLECULAR TRANSDUCER ACTIVITY | -2.11 |
| PROTEIN TYROSINE KINASE ACTIVITY | -2.12 |
| TRANSMEMBRANE RECEPTOR PROTEIN KINASE ACTIVITY | -2.15 |
| GROWTH FACTOR BINDING | -2.24 |
| EXTRACELLULAR MATRIX STRUCTURAL CONSTITUENT | -2.77 |

C.

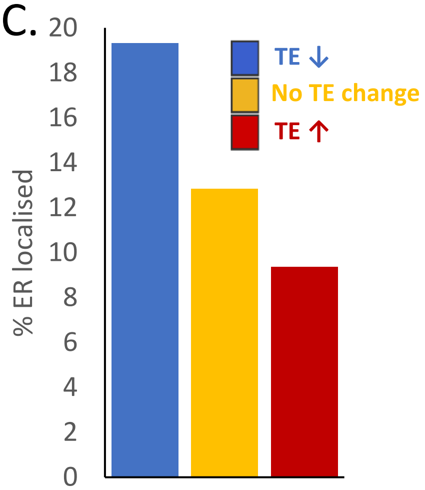


D.

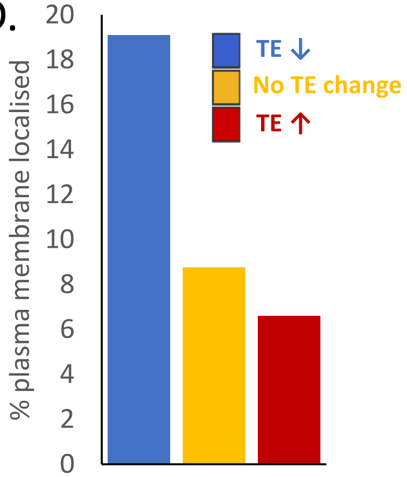

Fig. S8

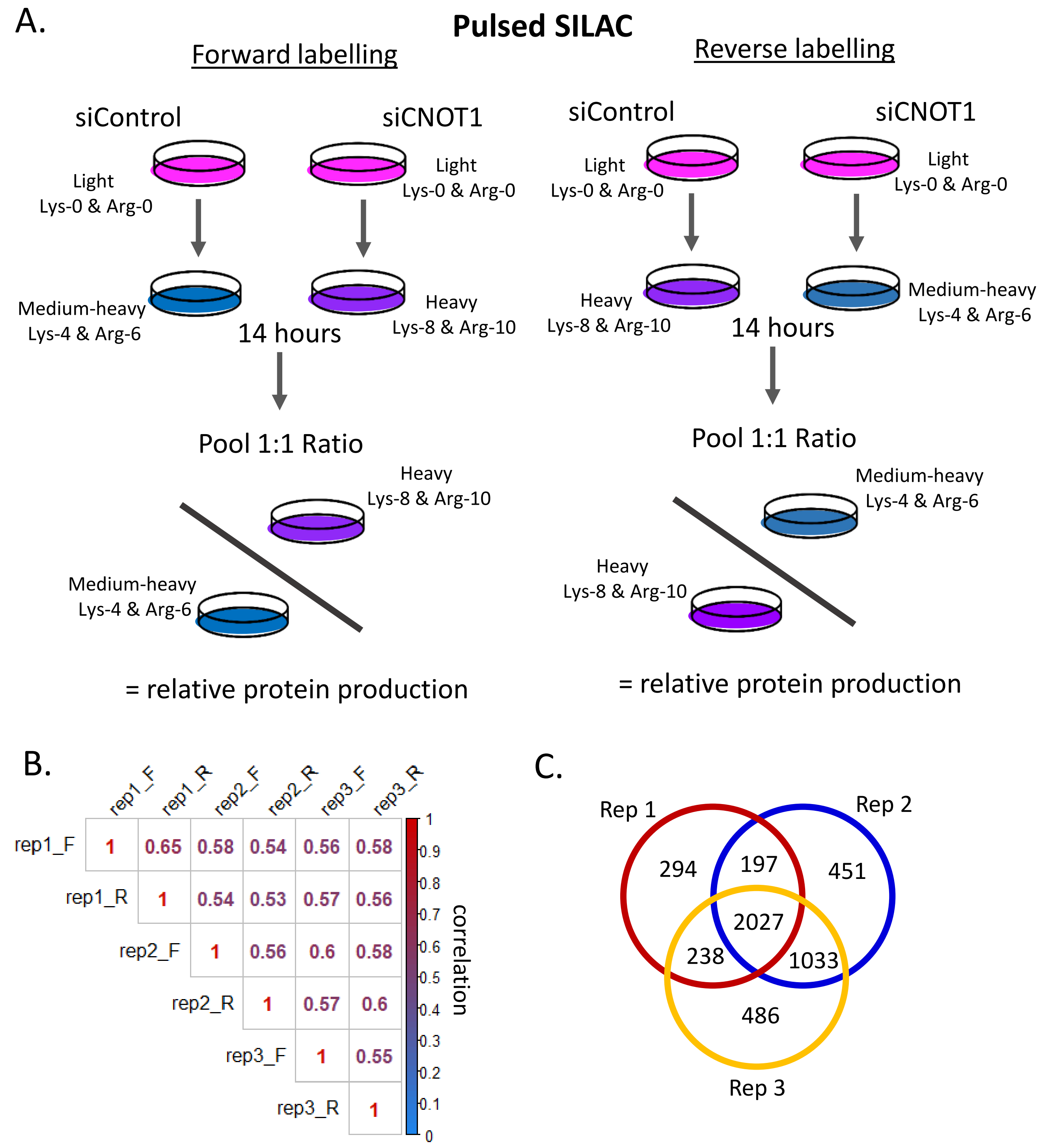
A.
Pulsed SILAC
Forward labelling
siControl
siCNOT1
Light
Lys-0 & Arg-0
Light
Lys-0 & Arg-0
Medium-heavy
Lys-4 & Arg-6
Heavy
Lys-8 & Arg-10
14 hours
Pool 1:1 Ratio
Heavy
Lys-8 & Arg-10
Medium-heavy
Lys-4 & Arg-6
= relative protein production
Reverse labelling
siControl
siCNOT1
Light
Lys-0 & Arg-0
Light
Lys-0 & Arg-0
Heavy
Lys-8 & Arg-10
Medium-heavy
Lys-4 & Arg-6
14 hours
Pool 1:1 Ratio
Medium-heavy
Lys-4 & Arg-6
Heavy
Lys-8 & Arg-10
= relative protein production
B.
rep1_F
rep1_R
rep2_F
rep2_R
rep3_F
rep3_R
rep1_F 1 0.65 0.58 0.54 0.56 0.58
rep1_R 1 0.54 0.53 0.57 0.56
rep2_F 1 0.56 0.6 0.58
rep2_R 1 0.57 0.6
rep3_F 1 0.55
rep3_R 1
correlation
1
0.9
0.8
0.7
0.6
0.5
0.4
0.3
0.2
0.1
0
C.
Rep 1
Rep 2
Rep 3
294
197
451
2027
238
1033
486

Fig. S9

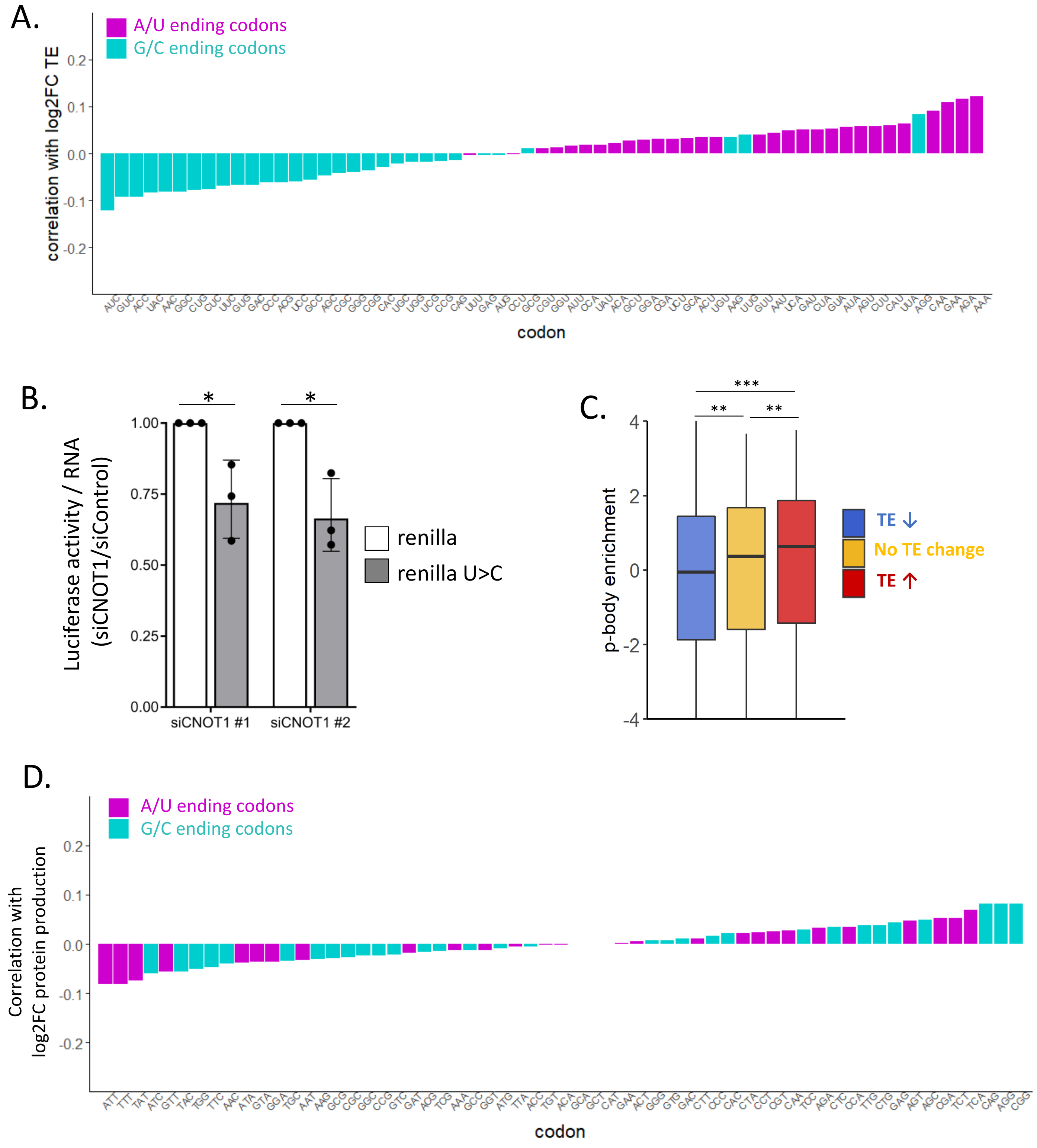
A.
A/U ending codons
G/C ending codons
correlation with log2FC TE
0.2
0.1
0.0
-0.1
-0.2
AUC GUC ACC UAC AAC GGC CUG CUC UUC GUG GAC CCC ACG UCC GCC AGC CGC GGG CGG CAC UGC UGG UCG CCG CAG UUU GAG AUG CCU GCG CGU GGU AUU CCA UAU ACA GCU GGA CGA UCU GCA ACU UGU AAG UUG GUU AAU UCA GAU CUA GUA AUA AGU CUU CAU UUA AGG CAA GAA AGA AAA
codon
B.
Luciferase activity / RNA
(siCNOT1/siControl)
1.00
0.75
0.50
0.25
0.00
*
*
siCNOT1 #1
siCNOT1 #2
renilla
renilla U>C
C.
p-body enrichment
4
2
0
-2
-4
***
**
**
TE ↓
No TE change
TE ↑
D.
A/U ending codons
G/C ending codons
Correlation with
log2FC protein production
0.2
0.1
0.0
-0.1
-0.2
ATT TTT TAT ATC GTT TAC TGG TTC AAC ATA GTA GGA TGC AAT AAG GCG CGC GGC CCG GTC GAT ACG TCG AAA GCC GGT ATG TTA ACC TGT ACA GCA GCT CAT GAA ACT GGG GTG GAC CTT CCC CAC CTA CCT CGT CAA TCC AGA CTC CCA TTG CTG GAG AGT AGC CGA TCT TCA CAG AGG CGG
codon

Fig. S10

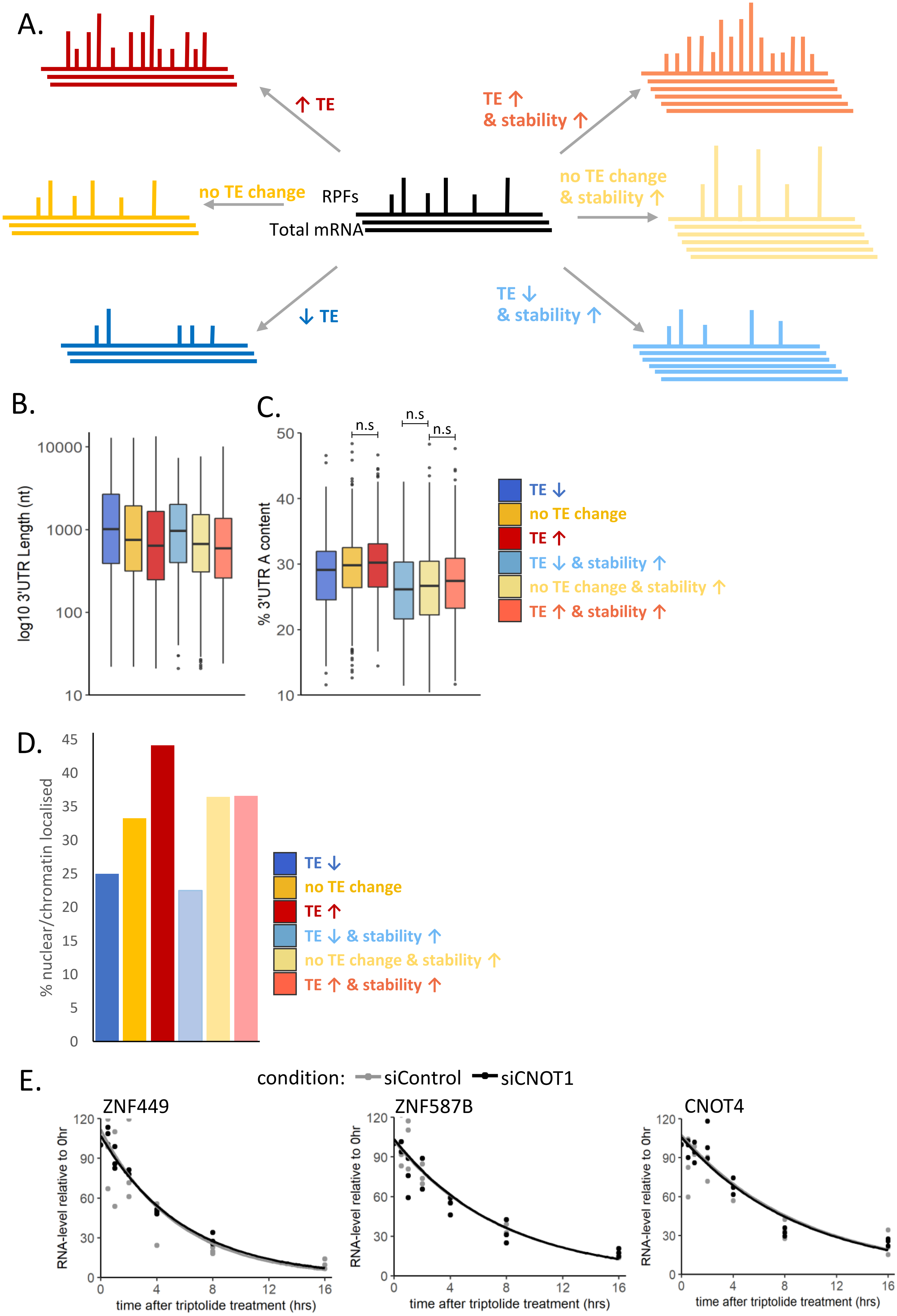

A.
↑ TE
TE ↑
& stability ↑
no TE change
RPFs
Total mRNA
no TE change
& stability ↑
↓ TE
TE ↓
& stability ↑
B.
log10 3'UTR Length (nt)
10000
1000
100
10
C.
% 3'UTR A content
50
40
30
20
10
n.s
n.s
n.s
TE ↓
no TE change
TE ↑
TE ↓ & stability ↑
no TE change & stability ↑
TE ↑ & stability ↑
D.
% nuclear/chromatin localised
45
40
35
30
25
20
15
10
5
0
TE ↓
no TE change
TE ↑
TE ↓ & stability ↑
no TE change & stability ↑
TE ↑ & stability ↑
E.
condition: siControl siCNOT1
ZNF449
ZNF587B
CNOT4
RNA-level relative to 0hr
120
90
60
30
0
0
4
8
12
16
time after triptolide treatment (hrs)

Fig. S11

A. Ribosome Protected Fragment (RPF)

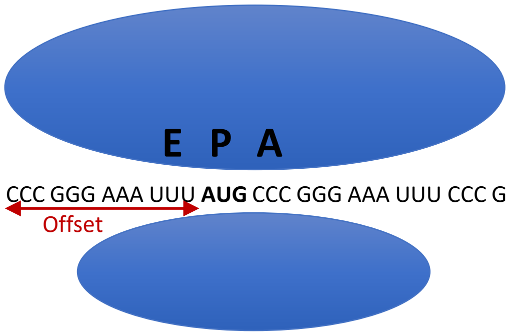


B.

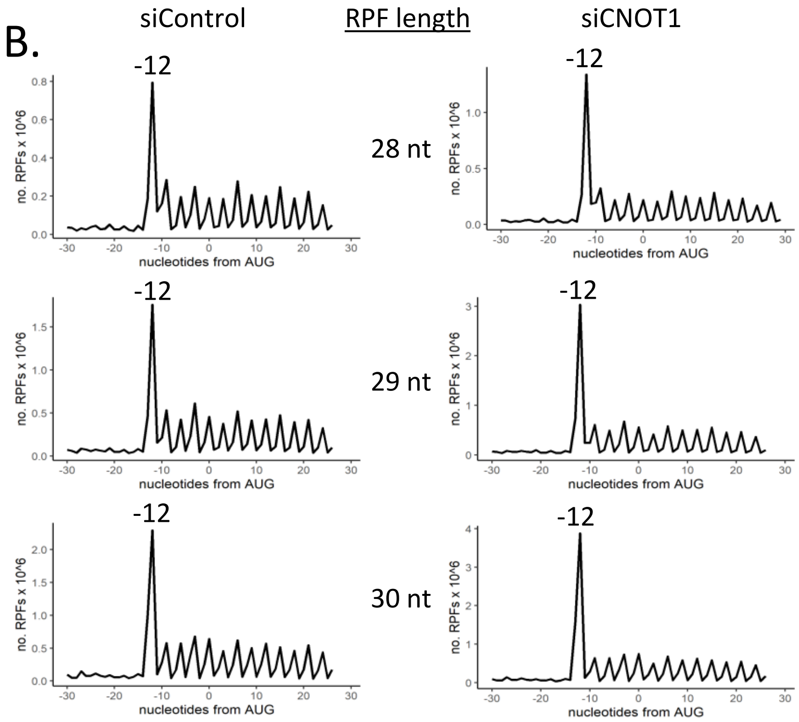
