## Supplemental Statistics for "Differential regulation of mRNA fate by the human Ccr4-Not complex is driven by CDS composition and mRNA localisation"

Additional File 2: Fig. S1 adjusted p-values for Main Fig. 4

### A. Protein production (Fig. 4B)

| No TE change | TE ↓ | TE ↑ & stability ↑ | No TE change & stability ↑ | TE ↓ & stability ↑ | |
|---|---|---|---|---|---|
| * | *** | ** | * | n.s | TE ↑ |
| | ** | *** | *** | * | No TE change |
| | | *** | *** | *** | TE ↓ |
| | | | * | ** | TE ↑ & stability ↑ |
| | | | | * | No TE change & stability ↑ |

*** p.adj < 0.001
** p.adj < 0.01
* p.adj < 0.1

### B. CDS length (Fig. 4D)

| No TE change | TE ↓ | TE ↑ & stability ↑ | No TE change & stability ↑ | TE ↓ & stability ↑ | |
|---|---|---|---|---|---|
| * | *** | *** | *** | * | TE ↑ |
| | *** | *** | *** | *** | No TE change |
| | | *** | *** | *** | TE ↓ |
| | | | ** | *** | TE ↑ & stability ↑ |
| | | | | ** | No TE change & stability ↑ |

*** p.adj < 0.001
** p.adj < 0.01
* p.adj < 0.1

### C. GC content (Fig. 4E)

| No TE change | TE ↓ | TE ↑ & stability ↑ | No TE change & stability ↑ | TE ↓ & stability ↑ | |
|---|---|---|---|---|---|
| n.s | ** | *** | *** | *** | TE ↑ |
| | ** | *** | *** | *** | No TE change |
| | | *** | *** | *** | TE ↓ |
| | | | n.s | n.s | TE ↑ & stability ↑ |
| | | | | ** | No TE change & stability ↑ |

*** p.adj < 0.001
** p.adj < 0.01
* p.adj < 0.1

### D. AG content (Fig. 4F)

| No TE change | TE ↓ | TE ↑ & stability ↑ | No TE change & stability ↑ | TE ↓ & stability ↑ | |
|---|---|---|---|---|---|
| ** | *** | *** | *** | *** | TE ↑ |
| | *** | *** | *** | *** | No TE change |
| | | n.s | * | *** | TE ↓ |
| | | | n.s | *** | TE ↑ & stability ↑ |
| | | | | *** | No TE change & stability ↑ |

*** p.adj < 0.001
** p.adj < 0.01
* p.adj < 0.1

Additional File 2: Fig. S2 adjusted p-values for Main Fig. 5C & Additional File 1: Fig. S10

A. Disorder promoting AAs (Fig. 5C)

| No TE change | TE ↓ | TE ↑ & stability ↑ | No TE change & stability ↑ | TE ↓ & stability ↑ | |
|---|---|---|---|---|---|
| *** | *** | *** | *** | *** | TE ↑ |
| | *** | *** | *** | *** | No TE change |
| | | n.s | n.s | n.s | TE ↓ |
| | | | n.s | n.s | TE ↑ & stability ↑ |
| | | | | n.s | No TE change & stability ↑ |

*** p.adj < 0.001
** p.adj < 0.01
* p.adj < 0.1

B. 3’UTR length (Fig. S10B)

| No TE change | TE ↓ | TE ↑ & stability ↑ | No TE change & stability ↑ | TE ↓ & stability ↑ | |
|---|---|---|---|---|---|
| * | *** | *** | *** | * | TE ↑ |
| | *** | *** | *** | *** | No TE change |
| | | *** | *** | *** | TE ↓ |
| | | | ** | *** | TE ↑ & stability ↑ |
| | | | | ** | No TE change & stability ↑ |

*** p.adj < 0.001
** p.adj < 0.01
* p.adj < 0.1

C. 3’UTR A content (Fig. S10C)

| No TE change | TE ↓ | TE ↑ & stability ↑ | No TE change & stability ↑ | TE ↓ & stability ↑ | |
|---|---|---|---|---|---|
| n.s | ** | *** | *** | *** | TE ↑ |
| | * | *** | *** | *** | No TE change |
| | | * | *** | *** | TE ↓ |
| | | | n.s | * | TE ↑ & stability ↑ |
| | | | | n.s | No TE change & stability ↑ |

*** p.adj < 0.001
** p.adj < 0.01
* p.adj < 0.1
